# Beyond traditional biodiversity fish monitoring: environmental DNA metabarcoding and simultaneous underwater visual census detect different sets of a complex fish community at a marine biodiversity hotspot

**DOI:** 10.1101/806729

**Authors:** Tania Valdivia-Carrillo, Axayácatl Rocha-Olivares, Héctor Reyes-Bonilla, José Francisco Domínguez-Contreras, Adrian Munguia-Vega

**Affiliations:** Laboratorio de Ecología Molecular, Departamento de Oceanografía Biológica, Centro de Investigación Científica y de Educación Superior de Ensenada (CICESE) Ensenada, Baja California, México; Laboratorio de Sistemas Arrecifales, Universidad Autónoma de Baja California Sur (UABCS) La Paz, Baja California Sur, Mexico; Conservation Genetics Laboratory, School of Natural Resources and the Environment, The University of Arizona, Tucson, AZ, USA; @ Lab Applied Genetics Research, La Paz, Baja California Sur, Mexico

**Keywords:** biodiversity monitoring, metabarcoding, environmental DNA, underwater visual census, reference database, fish community

## Abstract

Significant advances in the study of marine fish communities have been achieved with traditional monitoring methods and recently with novel genetic approaches. eDNA metabarcoding is one of them and a powerful tool for the study of biodiversity still in continuous development. Its applicability in marine ecology and conservation studies may be gauged by comparing its results with those of traditional methods. In the present investigation, we compare results from the underwater visual census (UVC) with eDNA metabarcoding (eDNA) carried out simultaneously in 24 rocky reef sites along the Gulf of California. We developed a two-PCR library preparation protocol followed by high throughput sequencing aimed at teleost fish. Our results show that both methods had different detection capabilities, and each registered different sets of fish taxa from rocky reefs, with some overlap. In particular, eDNA identified taxa from pelagic, demersal, and estuarine habitats beyond the rocky reef itself, suggesting differences in detection mainly attributed to the transport and permanence time of the eDNA in the ocean. Overlap in the detection with both methods increased with taxonomic level. We argue that substantial gaps in sequence reference databases for teleost are at the root of major discrepancies. Our results also confirm that PCR-based eDNA metabarcoding of seawater samples does not reflect patterns in abundance and biomass of species estimated from traditional methods. We discuss how to reconcile the results of eDNA metabarcoding and traditional methods in marine hotspots.

## INTRODUCTION

Detecting the presence and identity of organisms is a primary task in ecology, management, and in the conservation of biodiversity (Magurran, 2004). However, identifying the complete species components in a particular ecosystem is arduous, especially in the marine realm. In this sense, significant advances in the study of fish communities have been achieved traditionally through underwater visual censuses (UVC), catch records, traps, and baited remote underwater video, among other traditional methods (MacNeil et al., 2008). Still, all these monitoring practices are selective to particular groups of species, and show other sources of bias (Bacheler, Geraldi, Burton, Muñoz, & Kellison, 2017).

One of the most popular techniques (UVC), relies on multiple trained observers for recording presence, abundance, and length of individual fish within a fixed area. As a consequence, it is constrained by safety issues that limit the depth and duration of surveys, besides to logistical restrictions related to transporting SCUBA gear and personnel to remote locations. In addition, it is generally biased against wary, highly mobile organisms or small or cryptic species, and the detection could be adversely affected by local conditions (e.g., poor visibility and strong currents) (Bozec, Kulbicki, Laloë, Mou-Tham, & Gascuel, 2011; MacNeil et al., 2008).

Alternatively, environmental DNA (eDNA) metabarcoding has emerged as a powerful technique for monitoring marine eukaryotic biodiversity (Thomsen et al., 2012). It takes advantage of high-throughput sequencing of a conserved standard genomic region (*barcode*) (Hebert, Cywinska, Ball, & DeWaard, 2003), which is PCR-amplified from a complex environmental sample (Barnes & Turner, 2016; Taberlet, Coissac, Hajibabaei, & Rieseberg, 2012) in order to characterize communities of organisms.

Initial research indicates that eDNA metabarcoding is valuable for the detection of species (Boussarie et al., 2018; Günther, Knebelsberger, Neumann, Laakmann, & Martínez Arbizu, 2018; Stat et al., 2019) and, in this sense, there are advances in freshwater (Cilleros et al., 2018; Lecaudey, Schletterer, Kuzovlev, Hahn, & Weiss, 2019) and oceanic ecosystems (Kelly, Port, Yamahara, & Crowder, 2014; Miya et al., 2015; Yamamoto et al., 2017), mostly in temperate regions.

Despite the recent progress of marine eDNA metabarcoding studies, many challenges need to be resolved in order to successfully apply it as a regular method in biodiversity monitoring in natural environments (Deiner et al., 2017). In general, most of the complexity of this method is related to the ecology of eDNA and its effect on the detection and quantification of DNA molecules (Barnes & Turner, 2016). Particularly, in the marine ecosystem, eDNA transport and degradation time are of main importance e.g., some studies in coastal habitats suggest that marine eDNA detection is spatially variable and captures the presence of organisms at scales going from dozen meters (Jeunen et al., 2019; Port et al., 2016) to a few kilometers (O’Donnell et al., 2017a). These patterns are likely strongly influenced by local oceanographic features influencing eDNA movement at different spatiotemporal scales (Hansen, Bekkevold, Clausen, & Nielsen, 2018; Thomsen et al., 2012) and by the length of eDNA fragments (Jo et al., 2017), among other factors.

Further, to be fully useful as a monitoring method, eDNA techniques ideally should be able to provide quantitative estimates of organism-abundance to allow inferences regarding population size or status. For this, a direct relationship between amount of eDNA detected and organisms abundance or biomass is paramount (Spear, Groves, Williams, & Waits, 2015). This topic has been less explored in the ocean, nonetheless studies suggests that the correlation is weak (Elbrecht & Leese, 2015; Piñol, Mir, Gomez-Polo, & Agustí, 2015; Spear et al., 2015). Finally, another major limitation of this technique is the availability of complete reference databases of DNA sequences of morphologically identified species, which are required to assign eDNA reads to taxonomic entities (Andersen et al., 2019).

Here we evaluate and compare bony fish (Actinopterygii) community data derived from simultaneous traditional UVC and eDNA metabarcoding at rocky reefs from the Gulf of California (GOC, Mexico), a hyper-productive marine biodiversity hotspot located between temperate and tropical waters in the Eastern Pacific (Munguia-Vega et al., 2018). Specifically, our goals focused on: 1) Assessing the effect of the use of a complementary custom reference database (Custom DB) in the taxonomic assignment of eDNA reads, in contrast to only using NCBI-GenBank; 2) Comparing patterns of community composition and similarity obtained by UVC and eDNA; 3) Assessing the detection sensitivity of metabarcoding in a sample of known composition (mock community); and 4) Determining the relationship between the number of DNA reads obtained with eDNA and individual abundance and biomass obtained with UVC. To achieve this, we sampled 24 sites associated with rocky reefs and seamounts from the Central and Northern GOC and developed a Custom DB for some of the commercially important local reef species. By simultaneously applying both methods on different traditional biodiversity measures, our study allowed us to evaluate the current limitations of both approaches.

## MATERIALS AND METHODS

### Study area: Gulf of California

The GOC is among the world’s most productive marine regions (Roberts et al., 2002), and its ichthyofauna is composed of over 800 species of bony fishes (Actinopterygii) (Hastings, Findley, & Van Der Heiden, 2010; Robertson & Allen, 2019). In this semi-closed sea, intertidal and sub-tidal rocky-reefs are the main habitats that support biodiversity, as there are over 900 islands and islets (Munguia-Vega et al., 2018). Rocky reefs shelter at least 333 species of bony fish species (Actinopterygii: Teleostei), corresponding to 40% of the total teleost fauna of the GOC (Brusca et al., 2005; Thomson, Findley, & Kerstitch, 2000). Geographically, the GOC lies at the intersection of the temperate and tropical faunal regions of the Eastern Pacific, and its ichthyofauna represents a mixture from at least three provinces in addition to some globally distributed and endemic species (Robertson & Cramer, 2009). The GOC has been divided in a Northern, Central and Southern regions, based on biogeographic affinities, oceanography, bathymetry and environmental characteristics (Figure 1) (Brusca et al., 2005; Hastings et al., 2010; Walker, 1960).

**Figure 1.**
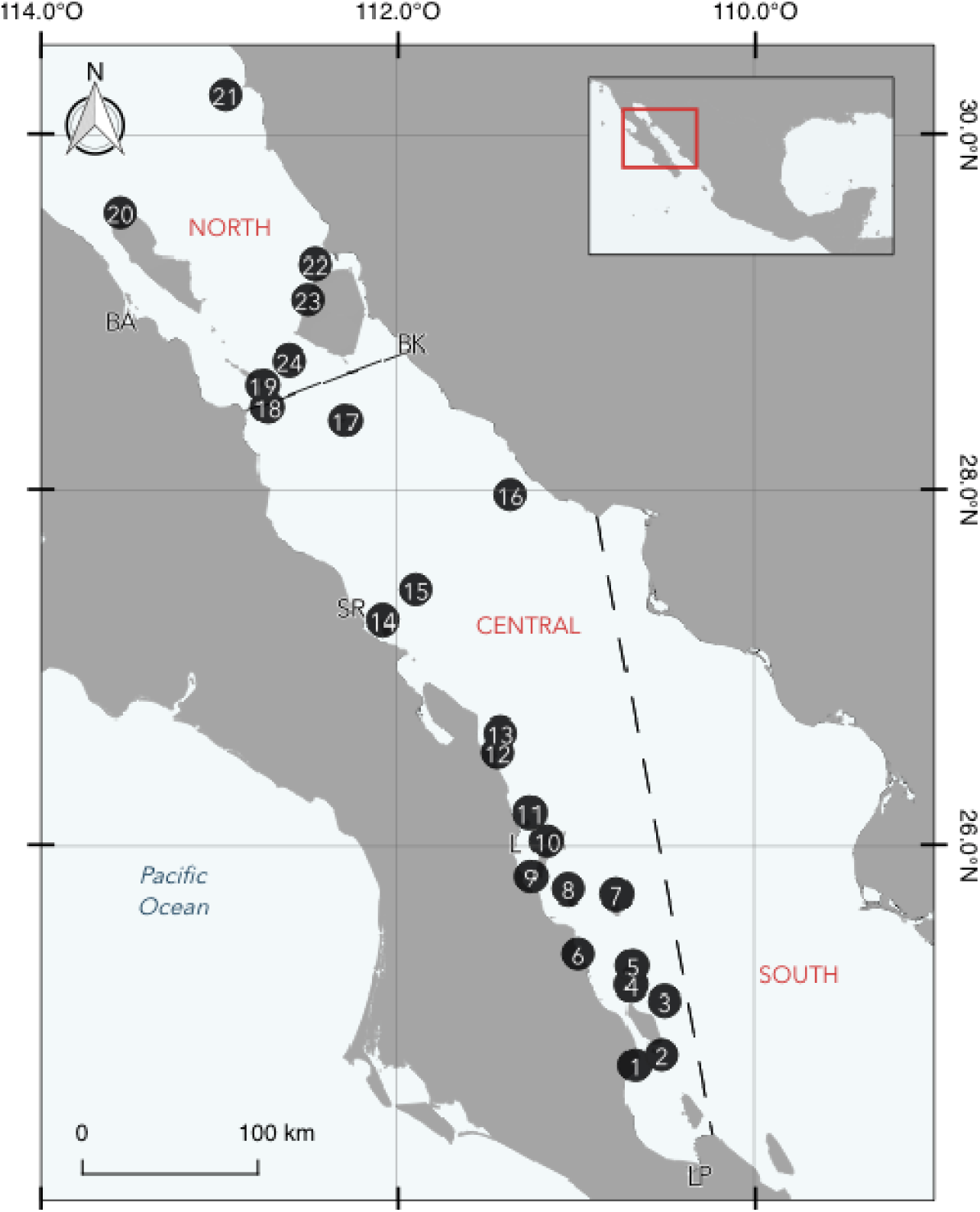
Sampling sites for UVC and seawater collection for eDNA metabarcoding across the Gulf of California (see Table 1 for site-specific details). Northern, Central, and South GOC regions are labeled. Principal towns are shown (BA: Bahia de los Angeles, SR: Santa Rosalia, L: Loreto, LP: La Paz, BK: Bahia de Kino).

### Underwater visual censuses (UVC) and seawater sampling for eDNA metabarcoding analysis

A scientific expedition was conducted from October 23 to November 15, 2016, onboard a 90-foot vessel to perform UVC and to collect seawater samples for eDNA metabarcoding at 24 rocky reefs across the GOC (Table 1 and Figure 1). Sampled sites included coastal regions, islands and seamounts. At each site, six divers used SCUBA gear and extensive fish identification expertise to identify and census conspicuous fishes (adults > 5 cm in total length) inside of 25 m-long transects, 4 m wide and 2 m high (i.e., 100 m^2^ surveyed for each transect). Transects were placed at different depths ranging from 10 to 25 m deep, parallel to the coastline or following the contour of the reefs (over seamounts), in order to cover as much habitat area as possible. For each fish species, divers visually estimated the total abundance as well as the total length (in intervals of 2.5 cm in fishes smaller than 15 cm, 5 cm if the measured organisms were between 15 - 40 cm, and 10 cm intervals for larger animals). From these data, individual size was converted to individual biomass using the species-specific length-weight relationship (biomass = a*Length in cm^b^/100), available in Fishbase (Froese & Pauly, 2019) or other sources (Balart et al., 2006; González Acosta, De La Cruz Agüero, & De La Cruz Agüero, 2004). Biomass estimates were computed from this relationship divided by 100.

**Table 1.** Sampling sites and associated metadata.

| Collection date | Site # | ID | Site Name | Lat | Long | Temp (°C) | Depth (m) | N transects |
| --- | --- | --- | --- | --- | --- | --- | --- | --- |
| 10/29/16 | 1 | POR | El Portugués | 24.749 | 110.674 | 28 | 6 | 14 |
| 10/29/16 | 2 | BSS | Bajo Seco Sur | 24.809 | 110.525 | 27 | 21 | 6 |
| 10/30/16 | 3 | ANI | Bajo Las Ánimas | 25.114 | 110.508 | 28 | 10 | 11 |
| 10/30/16 | 4 | SDI | Isla San Diego | 25.204 | 110.695 | 28 | 5 | 14 |
| 10/30/16 | 5 | SCR | Isla Santa Cruz | 25.313 | 110.689 | 28 | 11 | 10 |
| 10/31/16 | 6 | MAT | Isla San Mateo | 25.379 | 110.993 | 28 | 8 | 14 |
| 10/31/16 | 7 | CAT | Isla Catalana | 25.715 | 110.776 | 28 | 12 | 14 |
| 11/01/16 | 8 | MON | Isla Monserrat | 25.744 | 111.049 | 28 | 7 | 14 |
| 11/01/16 | 9 | DAN | Isla Danzante | 25.813 | 111.255 | 27 | 13 | 15 |
| 11/02/16 | 10 | CAR | Isla Carmen | 26.017 | 111.169 | 26 | 6 | 15 |
| 11/02/16 | 11 | COR | Isla Coronados | 26.174 | 111.262 | 27 | 9 | 10 |
| 11/03/16 | 12 | PUL | Punta Pulpito | 26.515 | 111.443 | 27 | 4 | 16 |
| 11/03/16 | 13 | ILD | Isla San Ildefonso | 26.625 | 111.427 | 27 | 9 | 18 |
| 11/04/16 | 14 | SMAR | Isla San Marcos | 27.260 | 112.089 | 25 | 8 | 16 |
| 11/04/16 | 15 | TOR | Isla Tortuga | 27.431 | 111.062 | 26 | 17 | 8 |
| 11/06/16 | 16 | NOL | Isla San Pedro Nolasco | 27.966 | 111.371 | 24 | 20 | 24 |
| 11/07/16 | 17 | PMA | Isla San Pedro Martir | 28.382 | 112.296 | 21 | 13 | 16 |
| 11/08/16 | 18 | FRA | Isla San Francisquito | 28.441 | 112.268 | 23 | 10 | 8 |
| 11/08/16 | 19 | LOR | Isla San Lorenzo | 28.585 | 112.759 | 23 | 13 | 8 |
| 11/09/16 | 20 | IA-I | Isla Ángel de la Guarda I | 29.555 | 113.559 | 24 | 11 | 16 |
| 11/11/16 | 21 | LOB | Puerto Lobos | 30.222 | 112.966 | 22 | 3 | 8 |
| 11/12/16 | 22 | PAT | Isla Pato | 29.266 | 112.464 | 19 | 9 | 8 |
| 11/12/16 | 23 | TIB | Isla Tiburón | 29.065 | 112.506 | 18 | 12 | 4 |
| 11/13/16 | 24 | EST | Isla San Esteban | 28.722 | 112.613 | 18 | 8 | 8 |

Simultaneous to the UVC, a different group of divers sampled 1 L of seawater using clean Nalgene™ Wide-Mouth HDPE Bottles (Thermo Scientific™). After seawater collection, bottles were closed underwater and remained closed until water filtration. In all cases, water was filtered immediately after collection using 0.44 μm hydrophilic nitrocellulose Millipore® filters placed in a Millipore® Sterifil® filtration system connected to a manual vacuum pump, used to accelerate the process. Each filter was removed from the filtration system, folded inwards, stored in 1.6 mL tubes (Neptune ®) with STE sterile buffer (100 mM NaCl, 1 mM EDTA, 5 mM Tris/HCl pH 7.5) and stored frozen (−20 ° C) in the laboratory. Between each sample, all the materials and bottles were washed with a 10% solution of chlorine (Clorox®) and rinsed repeatedly with abundant freshwater.

### Custom DB construction: primer design, PCR amplification, and sequencing

Partial or complete teleost mtDNA sequences for the 12S rRNA gene were obtained from NCBI for species present in the GOC, and aligned along with the mitogenome of *Cyprinus carpio* with the Muscle v3.8.31 plugin in Geneious Prime® version 2019.0.4. Primers Teleo12S_1322-R, Teleo12S_682-F, and Teleo12S_792-F (see Table **S1** in Supplementary material for details) were designed with the Primer3 implementation in Geneious Prime® in order to amplify 530 - 640 bp regions of the 12S rRNA gene, surrounding the 65 bp region amplified with “teleo” primers (Valentini et al., 2016). DNA from 67 morphologically identified fish species (1-8 specimens per species) was extracted with a salt protocol (Aljanabi & Martinez, 1997) and then used for PCR amplification with the newly designed primers. These specimens belong to commercial species that were collected in the GOC over the last decade as part of the ecosystem-based project PANGAS (Munguía-Vega et al., 2015) (Supplementary Table **S2**).

Amplifications were carried out on 25 μL reaction volumes containing 1x PCR Buffer Mg^+^ free, 1.5 mM MgCl_2_, 0.8 mM dNTP mix, 0.3 μM of each primer, 1.5 U of DNA Taq polymerase (Invitrogen), 1% BSA and 2 μL of DNA template. The thermal cycling profile was denaturation at 95ºC for 60 s; 30 cycles at 95ªC for 30 s, annealing at 61ºC for 15 s; and extension at 72ºC for 15 s; and a final extension at 72ºC for 5 min. Amplifications were visualized on 2% agarose gels stained with GelRed ® Nucleic Acid Gel Stain (Biotum). After PCR amplification, products were sequenced on an Applied Biosystems 3730XL DNA Analyzer at the University of Arizona Genetics Core Laboratory. Sequences were edited using Chromas Pro v1.6 and aligned using MUSCLE multiple alignment tools implemented in Mega6 (Tamura, Stecher, Peterson, Filipski, & Kumar, 2013). Annotation of the sequences was assisted using the MITOS web server (Bernt et al., 2013). The obtained sequences were used as a Custom DB for taxonomic assignment of sequence reads and registered in GenBank.

### eDNA extraction, library construction and high-throughput sequencing

Strict experimental protocols were implemented in the laboratory to avoid cross-contamination at all steps. eDNA extraction and library preparation took place in a separate hood with eDNA dedicated equipment, reagents and supplies, using sterilized filtered pipet tips. Each time the hood was used, ten minutes of UV light treatment was applied to sterilize the area. Negative controls were included in all lab processing steps.

eDNA extractions were completed with the Qiagen Blood and Tissue Kit (Qiagen, USA). For this, each filter was cut finely (approximately 1 mm^2^) with the help of sterile stainless-steel scissors and tweezers. Subsequently, the total STE buffer contained in each tube and the pieces of the filter were divided into two 1.6 mL tubes to proceed with DNA extraction. Extraction steps were carried out almost entirely following the supplier’s instructions, until the step previous to elution, in which the pieces of the filter were removed from the columns. Also, the final elution with AE buffer was performed in two separate final centrifugations: first adding 50 μL and then 100 μL. All samples were quantified by fluorescence with a HS assay kit for a Qubit 3.0 fluorometer (Invitrogen, CA, USA).

### Library construction and high-throughput sequencing

Library preparation involved two PCR steps. Primer sequences and PCR conditions are shown in Supplementary materials (Tables **S3** and **S4)**. In addition to environmental samples, a positive control (Mock community described below), and a negative control (DNAse free water) were incorporated during library preparation and sequenced.

#### Mock community

In order to test the detection sensitivity of metabarcoding in a sample of known composition, we designed a mock community experiment in which equal amounts (200 ng) of purified DNA from 22 reef fishes of the GOC (Table 2) were pooled and subsequently used as a positive control. This sample was incorporated to library construction process, and sequenced parallel to eDNA samples.

**Table 2.** Species of teleosts included in the mock community sample (n=22).

| Class | Order | Family | Genus | Species | Detection | Sequence<br>in<br>Custom<br>DB | Sequence in<br>NCBI |
| --- | --- | --- | --- | --- | --- | --- | --- |
| Actinopterygii | Perciformes | Balistidae | Balistes | *Balistes polylepis* | * | * |  |
| Actinopterygii | Tetraodontiformes | Balistidae | Sufflamen | *Sufflamen verres* | * | * |  |
| Actinopterygii | Perciformes | Carangidae | Seriola | *Seriola rivoliana* | * | * |  |
| Actinopterygii | Perciformes | Chaetodontidae | Johnrandalia | *Johnrandalia nigrirostris* | * | * | * |
| Actinopterygii | Perciformes | Cirrhitidae | Cirrhitus | *Cirrhitus rivulatus* | * | * |  |
| Actinopterygii | Perciformes | Haemulidae | Anisotremus | *Anisotremus interruptus* | * | * |  |
| Actinopterygii | Perciformes | Haemulidae | Haemulon | *Haemulon sexfasciatum* | * | * |  |
| Actinopterygii | Perciformes | Kyphosidae | Kyphosus | *Kyphosus elegans* |  | * | * |
| Actinopterygii | Perciformes | Labridae | Bodianus | *Bodianus diplotaenia* | * | * | * |
| Actinopterygii | Perciformes | Labridae | Thalassoma | *Thalassoma lucasanum* | * |  | * |
| Actinopterygii | Perciformes | Lutjanidae | Lutjanus | *Lutjanus argentiventris* | * | * |  |
| Actinopterygii | Perciformes | Lutjanidae | Lutjanus | *Lutjanus novemfasciatus* | * | * |  |
| Actinopterygii | Perciformes | Lutjanidae | Lutjanus | *Lutjanus peru* | * | * | * |
| Actinopterygii | Perciformes | Pomacanthidae | Holacanthus | *Holacanthus passer* | * | * |  |
| Actinopterygii | Perciformes | Pomacanthidae | Hoplopagrus | *Hoplopagrus guentheri* |  | * |  |
| Actinopterygii | Perciformes | Pomacentridae | Abudefduf | *Abudefduf troschelii* | * | * | * |
| Actinopterygii | Perciformes | Pomacentridae | Stegastes | *Stegastes rectifraenum* | * | * |  |
| Actinopterygii | Perciformes | Scaridae | Scarus | *Scarus ghobban* | * | * | * |
| Actinopterygii | Perciformes | Serranidae | Epinephelus | *Epinephelus labriformis* | * | * |  |
| Actinopterygii | Perciformes | Serranidae | Paralabrax | *Paralabrax aurogutatus* | * | * |  |
| Actinopterygii | Tetraodontiformes | Serranidae | Paranthias | *Paranthias colonus* | * | * | * |
| Actinopterygii | Perciformes | Serranidae | Semicossiphus | *Semicossiphus pulcher* | * |  | * |

#### First PCR step (PCR 1)

The first step was performed to amplify ~ 65 bp from the 12S rRNA gene using “teleo” primers reported previously (Valentini et al., 2016). Primers included a standard Illumina sequencing adapter, the gene-specific primer and an adapter used for the second PCR. To account for potential false negatives, an additive strategy was used, that offsets the variability in individual PCR replicates and thereby maximizes diversity detection. Tree independent PCR amplifications of each DNA sample were done, two of them with DNA from the first elution (50 μL) and one with eDNA from the second elution (100 μL).

#### Second PCR step (PCR 2)

The second PCR step was performed in order to incorporate dual molecular identifiers (MID) to individual samples on the same flow cell. Primer pairs used contain the appropriate 8-nt index sequence (Glenn et al., 2016), the adapter to bind to the first PCR, and the sequencing primer sites. Following to the second PCR, the three experimental replicates per sample were combined and purified using 1.8X volume of AmpureXP beads (Beckman and Coulter). After cleaning, each sample was quantified by fluorescence with the HS assay kit for Qubit. Finally, a total of 26 samples (24 sampling sites, one Mock community, and one negative control) were pooled into one 4 nM equimolar sample and sent to Genomic Services at Langebio-CINVESTAV. A single flow cell Illumina NextSeq 500 MID (35 Gb) v2 chemistry (2×150 bp paired-ends) was used for sequencing.

### Bioinformatic analyses

Bioinformatic analyses (Figure 2) were implemented with a Unix shell script, which incorporates command line tools as well as calls to third-party software. In the first step of the analysis, bcl2fastq v2.19 (Illumina) was used to de-multiplex indexed sequences. Then, Obitools v1.2.11 (Boyer, Mercier, Bonin, Taberlet, & Coissac, 2014) was employed for applying a robust primary filtering, which selects for high-quality, full-length sequences (detailed pipeline described in Supplementary material **S5**). Briefly, forward and reverse sequence reads were aligned and merged (*illuminapairedend)*, and only sequences with complete overlap were maintained for the next steps. Then, a quality filter was applied (*obigrep*) to select the reads with a minimum Phred-scaled base quality of 40 (precision of 99.99%). Sample/experiment assignation based on barcodes and primers was made (*ngsfilter*) to keep sequences with maximum two errors on primers (default parameter). A second filter was done (*obigrep*) to keep reads with length >59 and <66 bases, and to remove sequences with ambiguous nucleotides. A concatenation step was performed to obtain a file containing all the sequences in samples. Then, a de-replication step was used (*obiuniq*) to group identical reads. Here, the number of grouped reads was incorporated in the sequence header as an attribute (count) to simplify the process of comparing sequences against a database. Next, vsearch v2.7.0 was employed (Rognes, Flouri, Nichols, Quince, & Mahé, 2016) for chimera detection *de novo*, using the uchime_denovo algorithm. A step by step aggregation analysis was implemented in Swarm v2.2.2 (Mahé, Rognes, Quince, de Vargas, & Dunthorn, 2015) to cluster reads with a d=2 resolution. This value was selected according to Cawthorn, Steinman, & Witthuhn (2012) based on mean genetic divergence in 53 marine fish species in 43 genus and 23 families, where d = 2 corresponds to 97% identity threshold for OTU detection. Finally, singletons were removed after clustering, and OTUs/site table conversion was performed with the owi_recount_swarm R script (Wangensteen, 2019).

**Figure 2.**
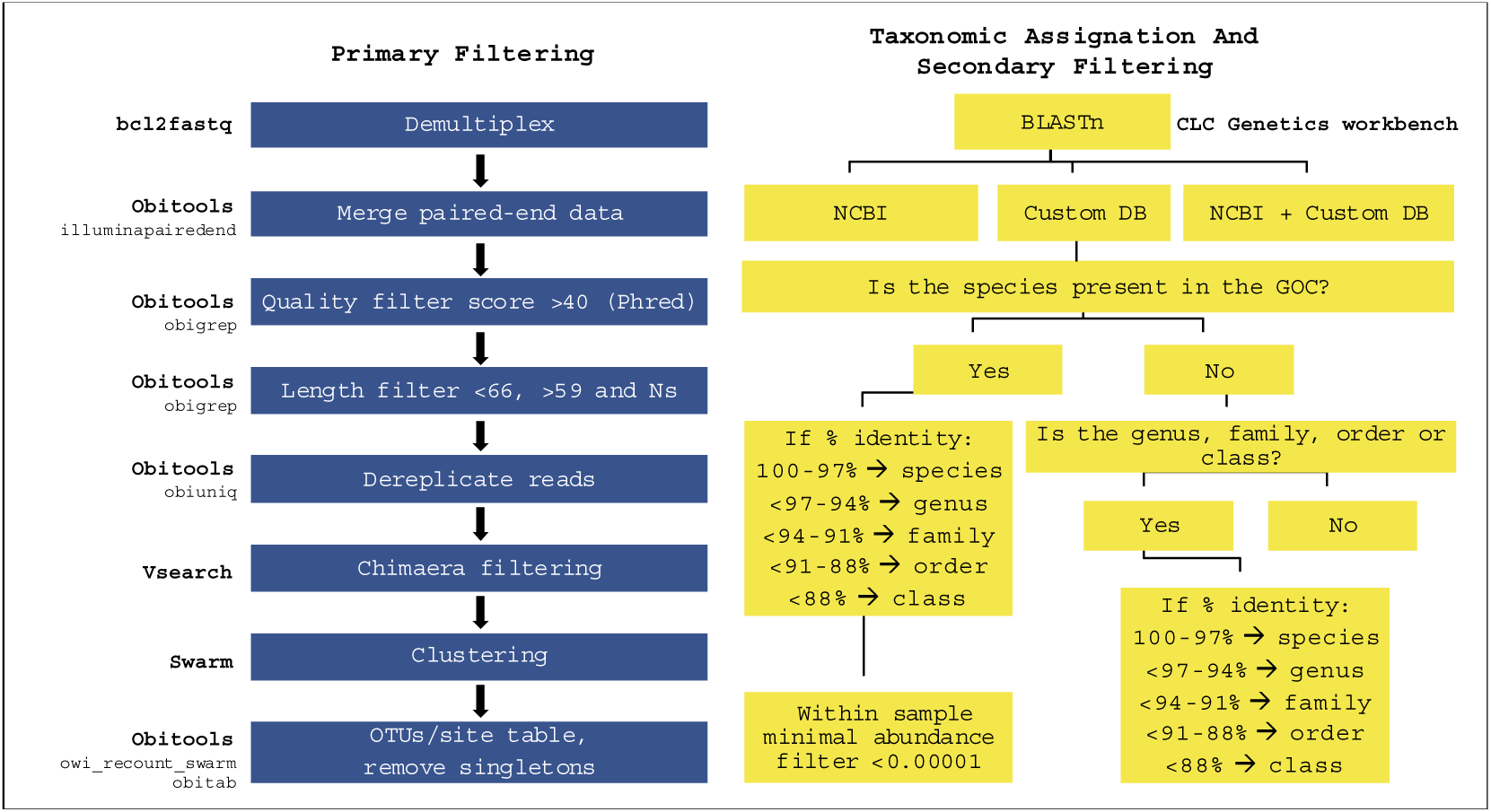
Bioinformatic workflows including primary filtering, taxonomic assignation and secondary filtering.

### OTUs taxonomic assignment

Taxonomic assignment of the OTUs was performed using three different approaches: 1) using NCBI-GenBank; 2) using our Custom DB of 67 commercial species, and 3) combining both (NCBI + Custom DB). To achieve this, a Basic Local Alignment Search Tool (Blast) implemented in CLC Genomics Workbench v9 (Qiagen) was used with a word count value of 30, expected score of 10 and e-value 1 e^−20^. OTUs without hits in the Blast search were excluded from taxonomic assignation. Conversely, in the resulting alignments, we used thresholds of the percent identity of the hits to assign taxonomy as follows: 100-97% of identity were assigned to species; < 97-94% to genus; < 94-91% to family; < 91-88% to order and < 88% to class (Cawthorn et al., 2012). Finally, taxonomic collapsing of distinct OTUs assigned to the same taxon was performed to obtain a final matrix containing the information of unique OTUs.

After these steps, secondary filtering was performed to avoid potential tag switching or false positives. We excluded OTUs with a minimal frequency within each sample <0.00001 (i.e., singletons in a locality) as supported by Bokulich and collaborators (2013) to improve diversity estimates from amplicon sequencing (Figure 2). To account for uneven sequencing depths, we rarefied each sample to the lowest observed value (59,014) using *rrarefy* function (*vegan*). Rarefaction curves were plotted using *rarecurve* function in the *vegan* package.

### Community-level ecological analyses

Lists of species/OTUs, genus, families, orders, and class detected with UVC and eDNA metabarcoding were generated. For community ecological analyses species/OTUs abundance data was transformed into presence/absence data to explore similarities among sampling methods in terms of community composition.

For each method, observed species/OTUs richness (S_*ob*s_) per locality was calculated with the *vegan* package in RStudio v1.2.1335, along with tests of statistical differences between methods (F-test and Welch two sample t-test). Likewise, Spearman’s correlation of S_*ob*s_ between methods was estimated in RStudio v1.2.1335. To assess sampling effort, S_*ob*s_ accumulation curves and expected species richness (S_*exp*_, Chao II) were plotted against sample sizes (liters filtered/transects surveyed) with Primer6 (K. R. Clarke & Gorley, 2006).

To summarize and compare site similarity patterns between methods a non-metric multidimensional scaling (nMDS) with 5000 iterations was executed, based on a presence/absence Jaccard similarity matrices in the *vegan* package in RStudio v1.1.447. Sampling locations were grouped in biogeographic regions of the GOC (Central and North, Figure 1). Statistical differences in OTU composition between regions were examined using permutational analysis of similarities (ANOSIM) with the same package. In order to asses if the faunistic composition derived from both methods were congruent, a Mantel test between UVC and eDNA Jaccard distance matrices between communities was performed using Spearman’s rs coefficient in Primer6 (K. R. Clarke & Gorley, 2006). Finally, correlations among eDNA sequence read abundance and UVC abundance and biomass per species/OTU was achieved using Spearman’s R coefficient in Past (Hammer, Harper, & Ryan, 2001).

## RESULTS

### Underwater visual censuses (UVC) species detection

UVC covered a total of 295 transects (mean = 12, SD = 4.5) in 24 sites from the GOC (Table 1). A total of 43,647 individual teleost fishes were observed, representing 97 species, 66 genera, 32 families, and 8 orders (Complete lists in **S6** Supplementary material).

### Custom DB for the ichthyofauna of the Gulf of California

We obtained 112 12S rRNA gene sequences from 67 species and 32 families with a mean length of 552 bp (SD = 65 bp) (NCBI-GenBank accessions in Supplementary material **S2**). Intra-specific variation was found in the species *Kyphosus elegans* (dmax 2.8%), *Lutjanus peru* (1.4%), *Mycteroperca rosacea* (1.4%), *Scomberomorus sierra* (1.3%) and *Thunnus albacares* (1.4%). Intra-generic variation ranged from 14.5 – 0%, and intra-family variation ranged from 42.9 – 0% (see Supplementary material **S7** for details).

### eDNA metabarcoding sequencing statistics

We obtained 5,429,682 paired-end reads with a length of 60-65 bases. Average sequencing depth per sampling site was 208,833 reads (range 121,385 - 323,473 reads). Rarefaction curves (Supplementary material **S8**) showed that a sequencing depth of 200,000 reads/site was sufficient to capture OTUs diversity among samples (Supplementary material **S9** for sequencing statistics per site during data filtering).

### OTUs identification in environmental samples, and taxonomic assignment

OTU clustering returned a total of 542 OTUs of the 12S rRNA barcode among the 26 samples. The success and resolution of taxonomic assignments varied depending on the reference database used (NCBI, Custom DB, NCBI + Custom DB, Table 3). Separately, the NCBI database and the Custom DB produced similar results in terms of the number of assigned OTUs (48% and 49% of 542 OTUs, respectively), whereas using both simultaneously increased the success identification by ~10% (60% of 542 OTUs). More than half of the OTUs assigned with NCBI were excluded from subsequent analyses to reduce the taxonomic uncertainty because they were assigned to taxa that have not been recorded in the GOC. In addition, the use of both databases simultaneously also outperformed the individual methods in terms of the taxonomic resolution at species and genus levels (Table 3). Thus, we used the NCBI + Custom DB taxonomic assignation results for further analyses. When OTUs could not be assigned to species or genus level, we named the OTU with a consecutive number, followed by the taxonomic level achieved according to our criteria, e.g., OTU_01 (Acanthuridae).

**Table 3.**
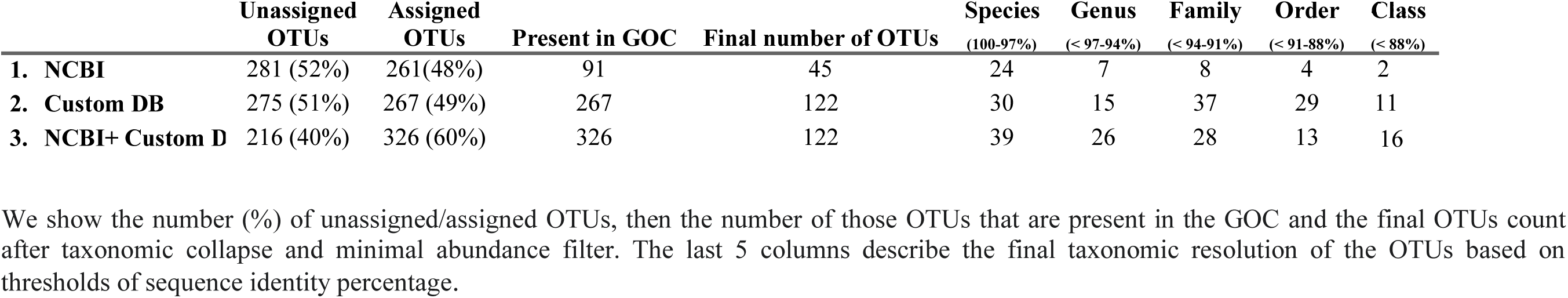
Summary of the taxonomic assignments of the 542 OTUs identified from the eDNA metabarcoding analysis employing three different reference databases: NCBI, Custom DB, NCBI + Custom DB (see methods for details).

After the taxonomic collapse, minimal abundance filtering and elimination of three OTUs classified as non-teleost (e.g. *Carcharhinus leucas*, *Gallus*, *Homo sapiens*), 122 unique Actinopterygii OTUs remained, of which 46.7% were taxonomically assigned above family level. The remaining 53.3% OTUs represent 117 species, 44 genera, 26 families, 7 orders and one class (Complete lists in **S10** Supplementary material).

### Mock community

Evaluation of the taxonomic identities in the mock community indicates that 20 out of the 22 species included (90.9%) were sequenced (Table 2). *Hoplopagrus guentheri* and *Kyphosus elegans* produced no identifiable reads in the mock community results despite having been included in the Custom DB. We detected one species that was not included in the mock community sample (*Mycteroperca rosacea*). For the negative control, only six reads were detected after quality filtering (two reads from *Mycteroperca rosacea*, two reads from *Thunnus albacares*, one read from *Paralabrax nebulifer*, and one read from *Lutjanus peru*). These results indicate negligible levels of foreign DNA contamination during DNA extraction, PCR, and library preparation.

### Community-level ecological analyses

Comparison of the taxonomic identities of the 194 species/OTUs identified in total using both monitoring approaches indicate that 12.9% (n = 25) were shared, 50% were only detected by eDNA (n = 97), and 37.1% were only registered with UVC (n = 72). The proportion of shared taxa between methods increased when higher taxonomic rankings were considered: 39.2% for genus (31/79), 52.6% for family (20/38), and 36.3% for order (4/11) (Figure 3).

**Figure 3.**
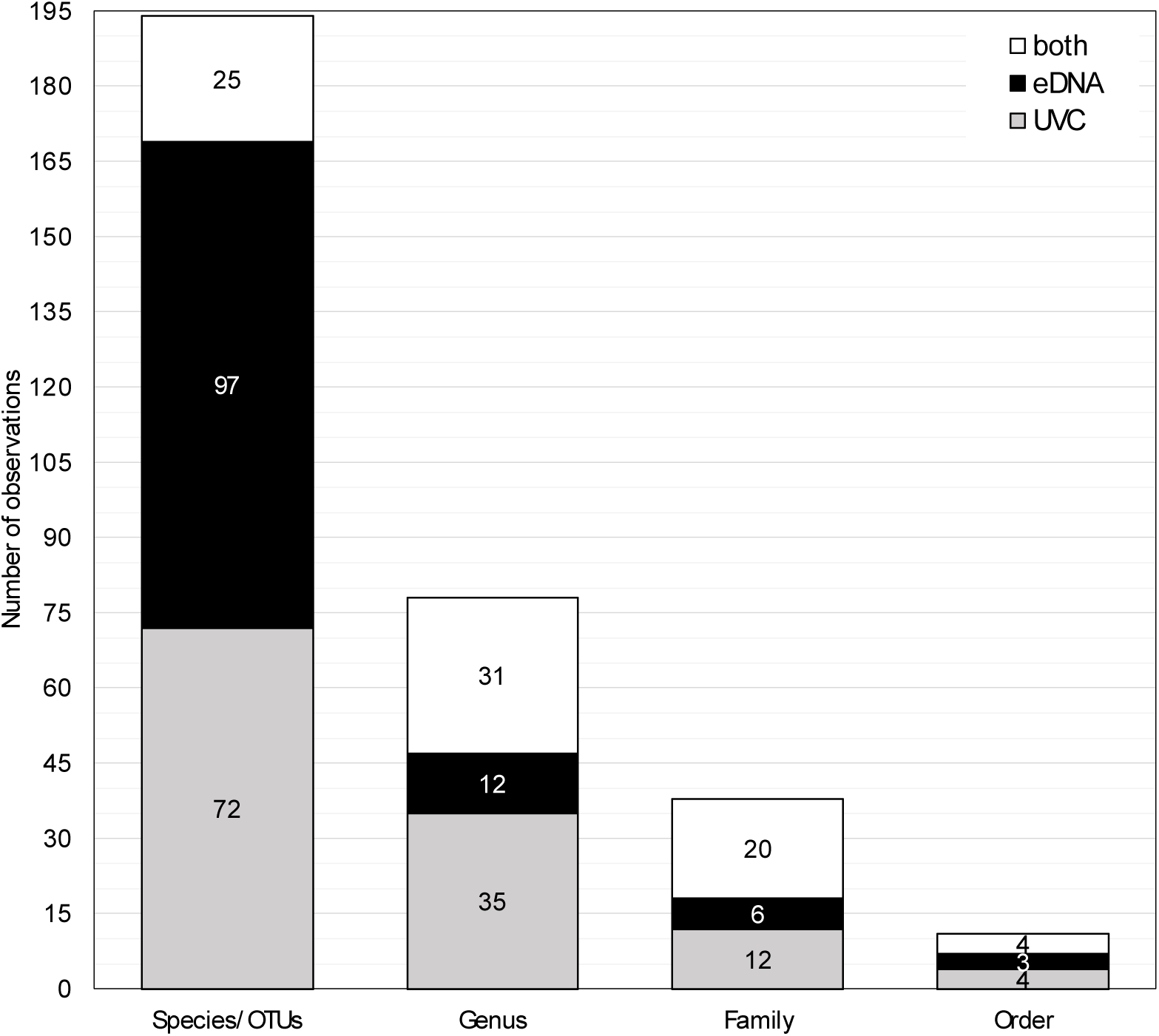
Number of observations at different taxonomic levels recorded by the monitoring methods: Underwater visual census (light gray; UVC), eDNA metabarcoding (black; eDNA), both (white). Numbers inside each diagram indicate the number Species/OTUs, Genus, Families and Orders.

Though UVC targeted only the class Actinopterygii, eDNA also recorded the class Chondrichthyes, Mammalia, and Aves, which were removed for the ecological analyses. Furthermore, while all the taxa registered in the UVC were reef-associated, eDNA metabarcoding detected additional taxa from surrounding pelagic, demersal and estuarine habitats. For example, we observed 20 families detected with both methods, six families that were only identified with eDNA and not UVC, including pelagic taxa (Istiophoridae, Scombridae, Clupeidae), demersal (Paralichthydae, Nomeidae) and estuarine (Mugilidae). Conversely, we observed that UVC detected 12 families that are reef-associated but that were not recovered by the eDNA methods (Figure 4). These results indicate that both techniques recorded distinct fish assemblages. For detailed lists of taxa at different taxonomic levels detected see **S11-S15** in Supplementary material.

**Figure 4.**
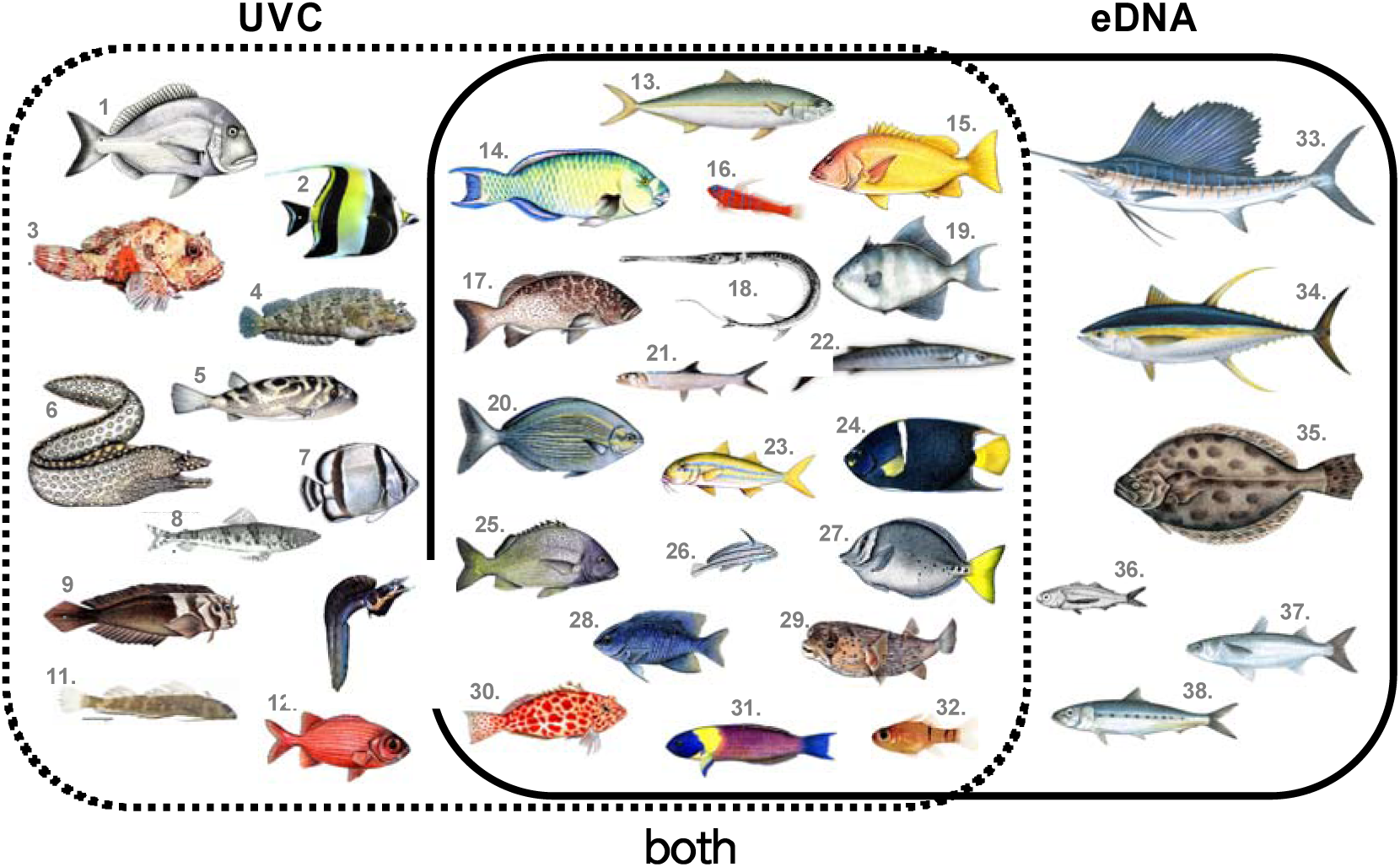
Families detected with each monitoring method. UVC: 1. Sparidae; 2. Zanclidae; 3. Scorpaenidae; 4.Labrisomidae; 5. Tetraodontidae; 6. Muraenidae; 7. Chaetodontidae; 8. Synodontidae; 9. Bleniidae; 10. Chaenopsidae; 11. Tripterygiidae; 12. Holocentridae. Both: 13. Carangidae; 14. Scaridae; 15. Lutjanidae; 16. Gobiidae; 17. Serranidae; 18. Fistulariidae; 19. Balistidae; 20. Kyphosidae; 21. Elopidae; 22. Sphyraenidae; 23. Mullidae; 24. Pomacanthidae; 25. Haemulidae; 26. Sciaenidae; 27. Acanthuridae; 28. Pomacentridae; 29. Diodontidae; 30. Cirrhitidae; 31. Labridae; 32. Apogonidae. eDNA: 33. Istiophoridae; 34. Scombridae; 35. Paralichthydae; 36. Nomeidae; 37. Mugilidae; 38. Clupeidae.

The mean number of observed (S_*obs*_) species/OTUs per site was significantly higher for eDNA than for UVC (t = −3.26, df = 39 p = 0.002; eDNA: 43 ± 6, range 33 – 54; UVC: 35 ± 9, range 18 – 50), and so was the variance (F = 2.45, df = 23, p = 0.03) (Figure 5 A). Species accumulation curves showed an incomplete saturation of species richness after 24 samples for both methods. The total number of expected species (S_*exp*_) estimated with Chao II for eDNA metabarcoding was also higher than for UVC (193 and 132, respectively) (Figure 5 B). The difference in the number of species detected between monitoring methods (S_*obs*_ eDNA/S_*obs*_ UVC) decreases significantly with increased sampling effort (number of transects surveyed), Spearman’s coefficient (R = −0.75, p=0.013). This indicates that the more transects surveyed, the number of species/OTU detected with the UVC becomes similar to that identified with the metabarcoding analysis of 1 liter of water (Figure 5 C). Spearman’s rs correlation of S_*obs*_ between methods was positive but marginally non-significant (R = 0.38, p = 0.066).

**Figure 5.**
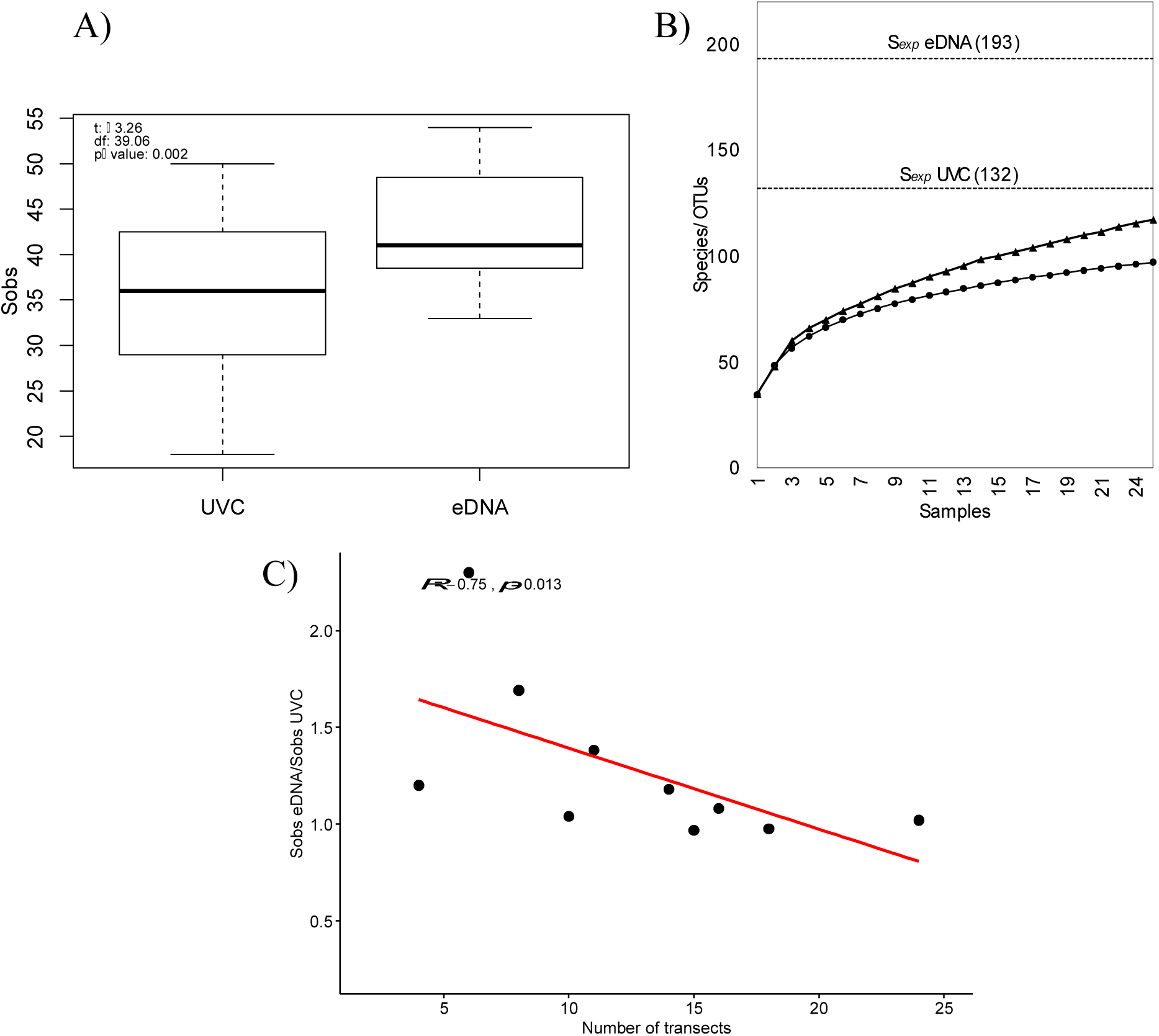
A) Boxplot of the mean observed number of species (S_*obs*_) for each sampling method for 24 sampling sites in the GOC. B) Observed species/OTUs rarefaction curves (incidence-based) for UVC (circles) and eDNA metabarcoding (triangles). The dashed line represents S_*exp*_ for the two methods. C) The ratio of species detection between monitoring methods (S_*obs*_ eDNA/ Sobs UVC) versus the number of transects surveyed, Spearman’s rs, and probability are shown. Linear regression in red and confidence intervals in gray.

Samples from Central and Northern biogeographic regions are discriminated by nMDS ordination analysis for UVC but not for eDNA metabarcoding (Figure 6). For UVC, a stress value of 0.127, indicated the presence of defined groups, in contrast to eDNA metabarcoding stress value 0.216 (Anosim R_UVC_ = 0.529, p = 0.006; Anosim R_eDNA_ = 0.111, p = 0.217). Mantel test among UVC and eDNA community distance matrices showed a non-significant correlation (R = −0.075, p = 0.72). On the other hand, mean S_*obs*_ from UVC and eDNA metabarcoding showed significant differences between North and Central regions (F_UVC_ = 13.18, df = 1, p = 0.001; F_eDNA_ = 11.28, df = 1, p = 0.002).

**Figure 6.**
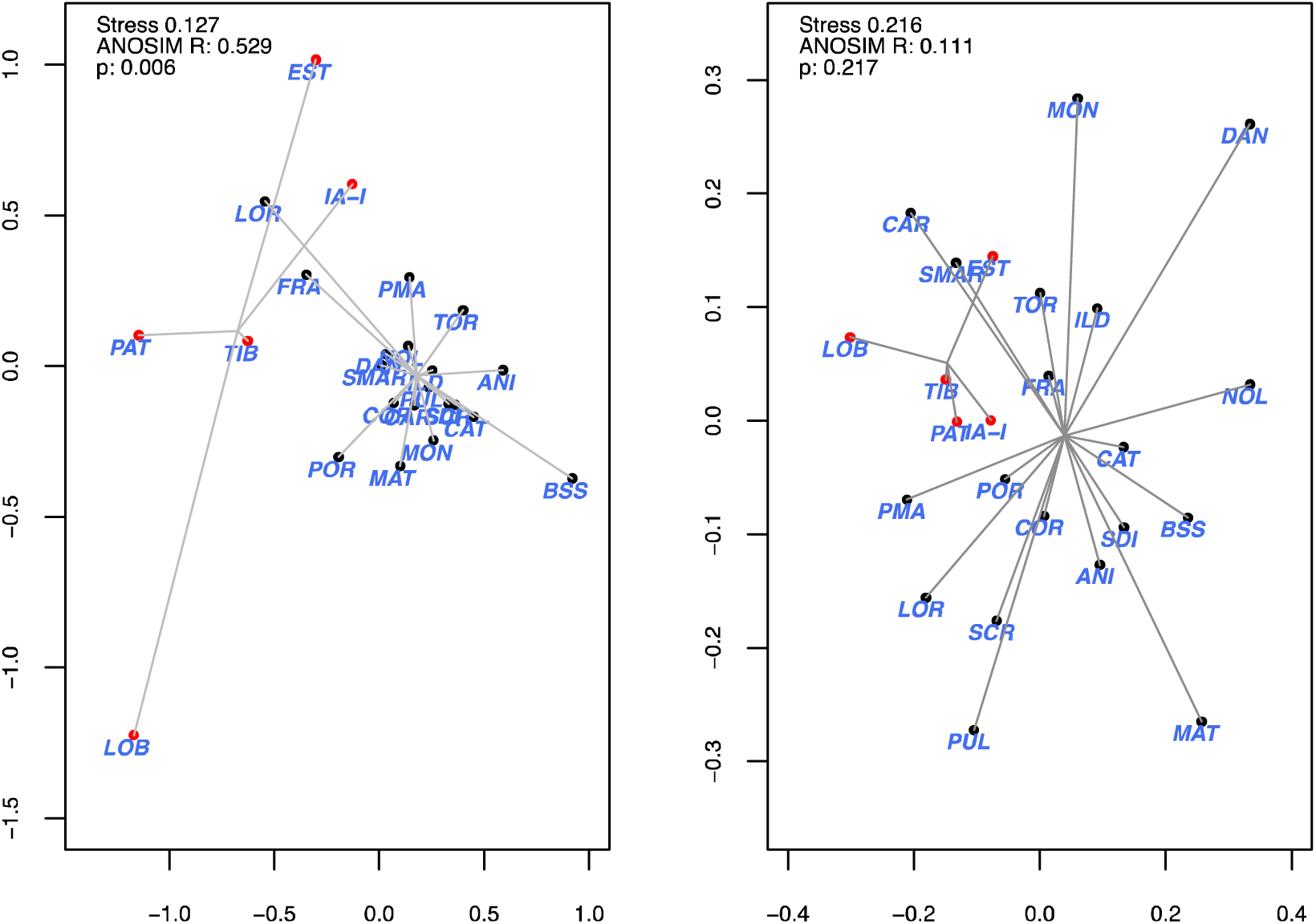
non-metric multidimensional scaling (nMDS) ordinations of UVC (A) and eDNA (B) samples using Jaccard similarity index.

**Figure 7.**
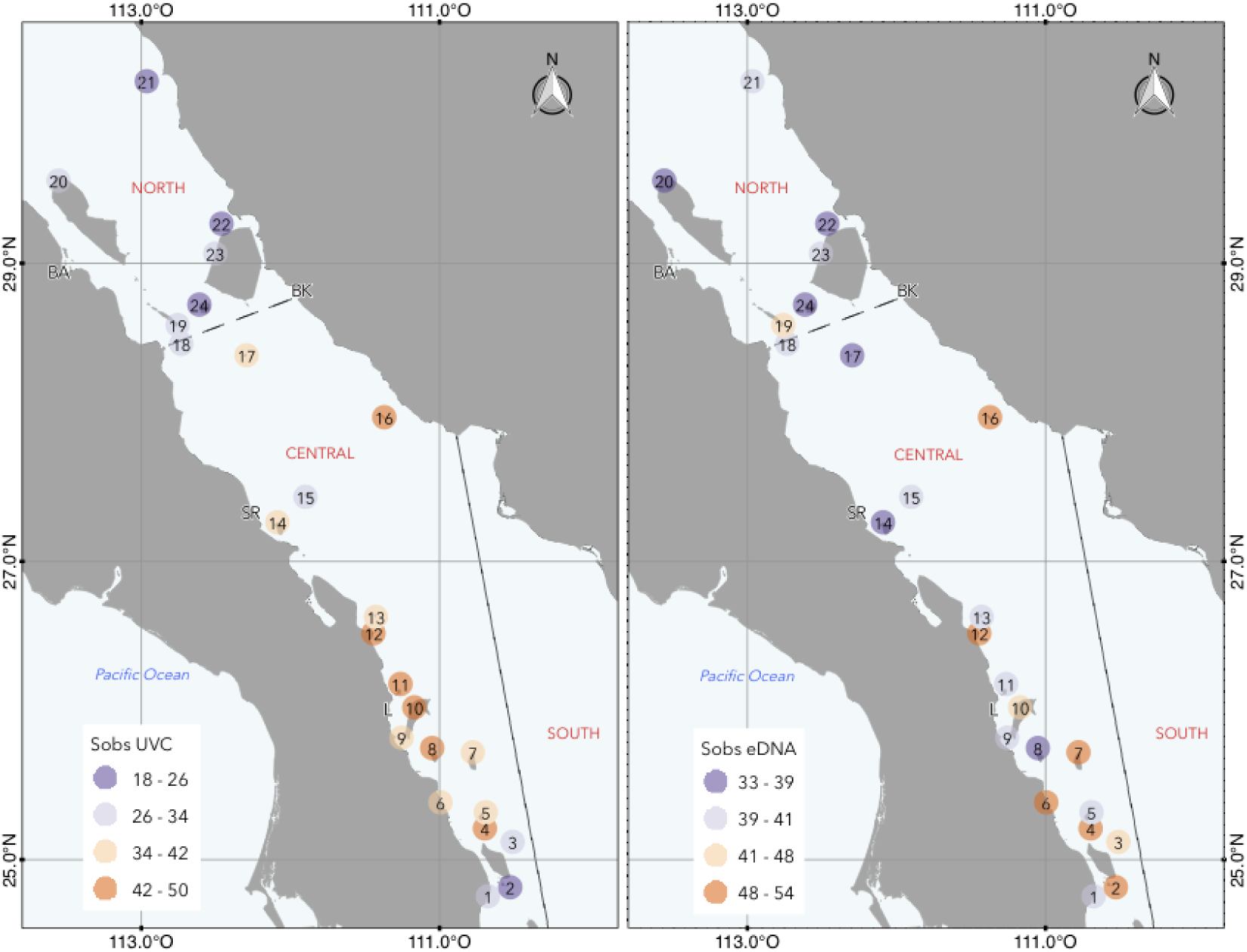
Geographical patterns of observed species richness with UVC (left) and eDNA metabarcoding (right).

Spearman’s correlation coefficients between eDNA number of reads for the 23 species shared by the two survey methods and their respective abundance and biomass estimated from UVC were not significant, except for the abundance of *Lutjanus argentiventris* (R_abundance_ = 0.525, p < 0.05) (**S16** Supplementary material).

## DISCUSSION

We compared two different monitoring methodologies focusing in evaluating teleost diversity in a marine hotspot: one traditional visual method widely used (UVC) and a novel genetic approach (eDNA metabarcoding) performed simultaneously. The results obtained here allow us to confirm the use of eDNA as an informative tool for the study of marine biological diversity. Our main findings indicate that UVC and eDNA have different detection capabilities when used as monitoring methods in high biodiversity marine ecosystems, with relatively low overlap. Moreover, our results show that some taxa were detected in UVC but not in the environmental samples, and vice versa. The molecular approach was able to detect, in addition to reef-associated species, others that are not persistent components of the reef ichthyofauna during their adult stage. Here, we discuss some of these outcomes concerning the strengths and limitations of this novel approach and provide recommendations for reconciling the results of eDNA metabarcoding and traditional approaches in future studies.

### UVC detection of species not found in eDNA metabarcoding

Several experimental and analytical factors drive the sensitivity of metabarcoding in detecting the presence of target species, which have to be identified in order to estimate their influence on measures of biodiversity (Calderón-Sanou, Münkemüller, Boyer, Zinger, & Thuiller, 2019; Furlan, Gleeson, Hardy, & Duncan, 2016; Kelly, Shelton, & Gallego, 2019). Some are related with technical aspects of data production such as primer affinity and PCR amplification preferences, PCR/sequencing errors and bias, and biological and technical replication; others to data analysis, such as reference databases completeness, criteria and parameters applied in bioinformatics pipelines, and ecological method of choice; and yet others even to inherent biological factors such as the evolutionary complexity of the community surveyed (Alberdi, Aizpurua, Gilbert, & Bohmann, 2017; Calderón-Sanou et al., 2019; Cristescu & Hebert, 2018; Kelly et al., 2019; Zinger et al., 2019).

In the present investigation, about 74% of the species observed in UVC were not detected in the eDNA samples. This translates to 44% at genera level, 31% at family and 36% at order level. The following are some of the possible factors that may be causing this incomplete detection of species. First, a source of taxonomic bias during the PCR amplification with "teleo" universal primers. We infer this possibility from the mock community experiment, in which two species (*Hoplopagrus guentheri* and *Kyphosus elegans*), which were included in the mock community DNA sample and for which a reference sequence was present in the reference DB, produced no detectable signal in the metabarcoding results. The first species (snapper) was recorded in the UVC surveys, yet it was also undetected in the environmental samples. Consequently, we recommend to estimate the taxonomic coverage of the barcode that will be used, and experimentally assess the possible primer mismatches with target DNA including representative taxonomic components of the fauna of interest (Kelly et al., 2019; Lecaudey et al., 2019). This can be achieved with *in silico* and *in vitro* PCR-amplifications (Bellemain et al., 2010; L. J. Clarke, Soubrier, Weyrich, & Cooper, 2014). Considering this possible source of bias, we also suggest that a second or third barcode (16S rRNA, COI) should be analyzed in parallel (Evans, Olds, Renshaw, et al., 2016) to the 12S rRNA barcode (Valentini et al., 2016). In addition, the use of blocking primers for unwanted DNA amplification, may also hinder amplification of target species with similar DNA sequences (Piñol et al., 2015; Port et al., 2016). Here, we used blocking primers during library preparation, to minimize amplification of human DNA (Valentini et al., 2016). Even though we did not test it, this could potentially inhibit amplification of some fish species. Yet, we suggest evaluating the co-blocking effect of the primer used for this purpose.

Additionally, concerning sequencing errors that may be interfering with species detection, stringent criteria were taken during all the bioinformatic pipeline analyses, including primary and secondary quality filters (Bokulich et al., 2013; Fonseca, 2018). Nevertheless, it may have resulted in a reduction in the number of OTUs detected and in their identification (Evans, Olds, Turner, et al., 2016).

In this regard, our results also point out to the need of a more thorough understanding of the dependency of eDNA metabarcoding on sampling effort. This is especially important in environmental samples taken from ecosystems which contain distinct species at very different concentrations (Piñol et al., 2015). Accordingly, we recommend increasing the number of biological and technical replicates to achieve a higher detection capability and compensate for other factors that may be biasing downwards the detection of some taxa (Cantera et al., 2019; Ficetola et al., 2015). A minimum of eight PCR replicates per sample for library preparation has been recently suggested to minimize PCR bias and maximize the detection probability of most species in a complex sample (Günther et al., 2018). Even so, a balance in the parameters used in PCR needs to be maintained to not substantially increase amplification errors during this step (Ficetola et al., 2015).

Concerning the replication on the field, and based on the incomplete saturation of the species accumulation curve of eDNA samples (Beentjes, Speksnijder, Schilthuizen, Hoogeveen, & van der Hoorn, 2019), we infer that the entire fish community was under sampled. In this sense, the sample volume/replicates required to adequately represent biological communities depends on taxon abundance, biomass, and the overall diversity of the community, and for that reason detection of species increases substantially when adding more replicated samples (Barnes & Turner, 2016; Shaw, Weyrich, & Cooper, 2016). Nevertheless, our results support the need for increasing biological replicates on the field particularly in high biodiversity regions.

Perhaps our main source of bias in species detection is attributable to the incompleteness of public DNA sequence databases, which restricts species taxonomic identification to available taxa. This limitation entails that species with missing DNA sequences can only be determined at best at higher taxonomic units (genera or families) (Cilleros et al., 2018). In light of the insufficient coverage of reference 12S rRNA gene sequences for the species of interest (only 5% of the ~800 teleost species from the GOC have DNA sequence data in NCBI-GenBank), we developed a Custom DB for 67 additional species, increasing the available genetic information to 13%. Regardless of the modest size of this reference database, the use of both NCBI + Custom DB increased from 24% to 39% the taxonomic identification at the species-level compared with using solely the information from NCBI. This gap in reference databases for the bony fishes of the GOC makes our results prone to Type II errors, referring to misidentification of queries without conspecifics in the database (Virgilio, Backeljau, Nevado, & De Meyer, 2010). Consequently, about half of the 122 OTUs could not be classified taxonomically. Hence, we emphasize that in order to fully exploit the potential in the detection power of eDNA metabarcoding, a vast effort is needed to improve taxonomic coverage of reference databases.

The Custom DB also showed us that, while the fixed divergence threshold used for taxonomic assignment seemed justified by some empirical data, the fish fauna from the GOC showed much larger variation within species, genera, and families (**S7** Supplementary material). The ichthyofauna of the GOC represents a mixture of evolutionary histories that converge in a single region (Hastings et al., 2010) and thus it is difficult to capture this phylogenetic variation using general thresholds of sequence divergence at a single locus. For example, some taxonomic families have very high sequence identity (Istiophoridae 98-100%) while others like Lutjanidae and Serranidae show much lower sequence identities ranging from 79.7-98.6% and 57.3 to 100%, respectively. Similarly, we observed that several species within the same family were indistinguishable from each other, e.g., *Rypticus bicolor*, *Epinephelus labriformis* and *Paranthis colonus* (Serranidae), and *Scarus ghobban* and S*. perrico*. This observed variation in the level of sequence divergence among closely related taxa has been reported previously for the 12S rRNA barcode (Cilleros et al., 2018; Valentini et al., 2016), which conflicts with the paradigm of a barcoding gap or non-overlapping intra- and inter-specific sequence divergence among closely related species to maximize taxonomic assignments (Collins et al., 2019; Cristescu & Hebert, 2018). However, it is unlikely that a single marker will meet all these criteria (Cristescu & Hebert, 2018), thus the use of multiple primers targeting the same taxonomic group could increase the taxonomic assignation breath and coverage (Alberdi et al., 2017; Lecaudey et al., 2019).

### Dark biodiversity: species detected in eDNA but missing from UVC

We found that 79% of the species/OTUs detected with eDNA were not registered during UVC, which translates to 29% at genera level, 23% at family level and 42% at order level. This visual non-detection might be in part due to observation limitations related to animal elusiveness, rareness, movement, density, size, habits, etc. (Bozec et al., 2011; Yamamoto et al., 2017). Also, factors related with eDNA ecology can be playing an important role (Barnes & Turner, 2016). For instance, the transport and permanence time of DNA molecules on the environment that allows molecules to move from adjacent locations to the sampled sites.

Our results indicate that all species detected during UVC were reef-associated, while eDNA registered some species that are not permanent residents of rocky reefs ecosystems, even if they are residents of the GOC. For example, this includes open-water pelagic species (*Cubiceps*, *Istiophorus*, *Katsuwomis*, *Sardinops*, *Sectator*, *Thunnus*), demersal fishes that are generally found at depths over 30 m (*Paralichthys*, *Hyporthodus*), and taxa associated with soft bottoms and estuaries (*Cynoscion*, *Mugil*, *Atractosion*).

From the technical and methodological point of view, precautions were taken during sampling, filtering, and during laboratory protocols to avoid contamination. Also, negative controls showed negligible number of reads for extraneous species. Therefore, our results suggest that eDNA metabarcoding integrate larger spatial and temporal scales in contrast with UVC. This represents a challenge in our understanding of complex processes that produce, transport and degrade eDNA in the environment. As suggested elsewhere, these processes are influenced by the physiology and space used by organisms as well as by oceanographic characteristics, suggesting that these factors ought to be incorporated into eDNA studies (Goldberg et al., 2016; O’Donnell et al., 2017a). In this sense, the persistence time of suspended eDNA can be highly variable (from 1 to 58 days; Hansen et al., 2018), although estimates vary by site, species and physicochemical conditions. Also, a recent study that used a particle tracking oceanographic model found that fish DNA could have originated 40 km south of the sampling site considering a decay time of 4 days (Andruszkiewicz et al., 2019).

These two characteristics imply that marine eDNA in the GOC could be transported as far as dozens of km, depending on local oceanographic conditions, before being detected (Marinone, 2012). Consequently, it would be useful to take into account oceanographic connectivity features during the period of sampling. In this context, a three-dimensional oceanographic model has shown that in seven days passive particles may be transported an average of 21 to 74 km in the northern GOC (Soria et al., 2012). Strong asymmetric seasonal currents, characteristic of the GOC, could transport eDNA from distant upstream sources, which may drastically shift north and south between warm and cold periods of the year (Munguia-Vega et al. 2018), highlighting marine eDNA results need to be interpreted in a temporally-explicit context.

Transport of eDNA and homogenization via oceanic currents may also explain why the nMDS from eDNA did not discriminate between North and Central biogeographic regions, whereas UVC did. First, this regionalization is defined by the geographic distribution of adult populations of reef fishes. In contrast, the species assemblages detected with eDNA incorporate the signal of additional species with broader distributions along their entire life-cycle, whose DNA may have been transported by mesoscale processes such as eddies or tidal exchange. Second, the uncertainty in species-level identification that underestimates sample species diversity could obscure the identification of biogeographic community patterns. The mixing of ecotones and communities caused by oceanographic features in marine eDNA studies agrees with other previous studies (Jeunen et al., 2019; O’Donnell et al., 2017b). Nevertheless, both the mean values and geographical patterns of S_*obs*_ of UVC and eDNA corroborate that the central region between Loreto and La Paz is a hotspot of species richness, diversity and abundance in contrast to the Northern GOC (Morzaria-Luna et al., 2018; Olivier et al., 2018). The spatial patterns revealed by eDNA need to be further investigated under the scrutiny of a more complete taxonomic assignation of the metabarcoding OTUs.

### Future directions for the application of eDNA metabarcoding in studies of marine biodiversity

eDNA metabarcoding holds a great potential for advancing the monitoring of marine biodiversity as it is an informative tool compared with traditional visual methods. Nevertheless, before the sources and fate of eDNA in the ocean are completely understood, comparative studies with traditional biodiversity monitoring methods will be key in establishing their limitations and potential applications for biodiversity monitoring. In this regard, a significant caveat in the use of PCR-based methods, such as eDNA, is their limitation as quantitative proxies of organism abundance, which restricts their use as presence/absence indicators.

The potential application of these techniques for the study of biodiversity is encouraging and is growing at an accelerated pace. In the marine environment, they are particularly valuable in areas that are logistically challenging for other survey methods, such deep mesophotic reefs between 50-200 m or bathypelagic ecosystems below 200 m. They have also been used in the study of fishery resources and in aspects related with biodiversity evaluations in marine reserves (Cristescu & Hebert, 2018; Evans & Lamberti, 2018).

Based on our study, we recognize increasing the number and quality of reference DNA sequences of marine fauna in public databases as a top priority. This effort requires investment and development of research groups including taxonomic experts and scientific collections that are focused on the development and standardization of field and laboratory methodologies and data analyses. Once these improvements are achieved, eDNA metabarcoding will allow for more efficient and cost-effective biodiversity monitoring. Increased automation will also make possible more frequent or continuous sampling to understand almost real-time changes in key species related to anthropogenic activities, including species of commercial or conservation value.

## ACKNOWLEDGEMENTS

We thank the support from Consejo Nacional de Ciencia y Tecnología, México (institutional fund CONACyT Fronteras de la Ciencia 292/2016), The University of Arizona—CONACYT CAZMEX consortium, Comunidad y Biodiversidad A.C., The Nature Conservancy Mexico, Pronatura Noroeste A.C. and Ecology Project International - La Paz. AMV received support from The David and Lucile Packard Foundation via the PANGAS Science Coordination. We received a grant from Langebio - Biotech del Norte for the Illumina Sequencing. This paper is part of the PhD research of TVC in the Marine Ecology postgraduate program at CICESE. She is also the recipient of a PhD scholarship granted by CONACyT (CVU 290839). We thank Manuel Alejandro Olán González, Damien Olivier, Jaime de la Toba, Omar Valencia and José Torres for helping with the UVC, Eldridge Wisely for helping with seawater sampling and filtering and Eric Ramírez for DNA extractions of the reference database. We thank the invaluable logistical support in the field from the crew of the DV Quino el Guardian, and also appreciate the drawings of fish from the Gulf of California originally made by Juan Chuy and provided by Sociedad de Historia Natural Niparaja A.C. The authors have no conflict of interest to declare.

## DATA ACCESSIBILITY

GenBank Bioproject PRJNA576668.

## AUTHOR CONTRIBUTIONS

TVC, AMV, HRB, ARO designed research; TVC, AMV performed research; TVC, AMV, HRB, ARO contributed new reagents or analytical tools; TVC, AMV, JFDC analyzed data: TVC, AMV, ARO, HRB wrote the paper.

