## Supplementary material for "Beyond traditional biodiversity fish monitoring: environmental DNA metabarcoding and simultaneous underwater visual census detect different sets of a complex fish community at a marine biodiversity hotspot"

**Table of Contents:**

| S1. Primer pairs for 12S rRNA amplification for Custom DB. | Page 2 |
| --- | --- |
| S2. Species included in the Custom DB and GenBank accession number. | Pages 3-4 |
| S3. Primers for library construction. | Page 5 |
| S4. Details for PCR 1 and 2 in library preparation. | Page 6 |
| S5. Pipeline for 12S rRNA metabarcoding bioinformatic analysis (OBITOOLS, VSEARCH, SWARM, R). | Page 7 |
| S6. Taxonomic list of Actinopterygii detected from UVC. | Page 8-10 |
| S7. Identity % percentage estimated at intra-genus and intra-family level with the 12S rRNA barcode. | Page 11 |
| S8. Rarefaction curves for eDNA metabarcoding sequence reads. | Page 12 |
| S9. Reads counts per bioinformatic step in primary filtering. | Page 13 |
| S10. Taxonomic list of Actinopterygii detected from eDNA metabarcoding**.** | Page 14-17 |
| S11. Species/OTU detection from UVC and eDNA metabarcoding. | Page 18-22 |
| S12. Genus detection from UVC and eDNA metabarcoding. | Page 22-24 |
| S13. Family detection from UVC and eDNA metabarcoding. | Page 24-25 |
| S14. Order detection from UVC and eDNA metabarcoding. | Pages 25 |
| S15. Class detection from UVC and eDNA metabarcoding. | Page 25 |
| S16. Spearman correlation between read abundance and organism abundance and biomass, for the shared species detected with UVC and eDNA metabarcoding in 24 sites of the GOC. | Page 26 |

**Table S1. Primer pairs for 12S rRNA amplification for Custom DB.**

| **Pimer** | **Name** | **Amplicon length**  **(bp)** | **Sequence (5´-3´)** | **Length** | **%GC** | **Hairpin**  **Tm** | **Pair Dimer**  **Tm** | **Self-Dimer**  **Tm** | **Tm** |
| --- | --- | --- | --- | --- | --- | --- | --- | --- | --- |
| **1,322 R** | Teleo12S_1322-R | 640 | CTTTCAGCTTTCCCTTGCGG | 20 | 55 | None | None | None | 59.8 |
| **682 F** | Teleo12S_682-F |  | CGTTCAACCTCACCCTTCCT | 20 | 55 | None | None | None | 59.6 |
| **792 F** | Teleo12S_792-F | 530 | CAGGTCGAGGTGTAGCGYATG | 21 | 60 | None | None | 11.5 | 60.3 - 63.4 |

#### Table S2. Species included in the Custom DB and GenBank accession number.

| **RefDB** | **Species** | **GenBank Accession number** |
| --- | --- | --- |
| **1** | *Abudefduf troschelli* | MK902806, MK902816 |
| **2** | *Acanthocybium solandri* | MK902889 |
| **3** | *Acanthurus xanthopterus* | MK902826 |
| **4** | *Anisotremus interruptus* | MK902835, MK902845, MK902894 |
| **5** | *Atractoscion nobilis* | MK902914 |
| **6** | *Balistes polylepis* | MK902859 |
| **7** | *Bodianus diplotaenia* | MK902867 |
| **8** | *Chaetodon humeralis* | MK902807, MK902817 |
| **9** | *Chanos chanos* | MK902919 |
| **10** | *Cirrhitus rivulatus* | MK902836 |
| **11** | *Coryphaena hippurus* | MK902902, MK902913 |
| **12** | *Cynoscion reticulatus* | MK902846 |
| **13** | *Cynoscion xanthulus* | MK902915 |
| **14** | *Diapterus brevirostris* | MK902906 |
| **15** | *Elacatinus puncticulatus* | MK902879, MK902886 |
| **16** | *Epinephelus acanthistius* | MK902884 |
| **17** | *Epinephelus labriformis* | MK902907 |
| **18** | *Gnathanodon speciosus* | MK902899 |
| **19** | *Haemulon sexfasciatum* | MK902852, MK902860 |
| **20** | *Haemulopsis leuciscus* | MK902868 |
| **21** | *Holacanthus passer* | MK902808, MK902818 |
| **22** | *Hoplopagrus guentheri* | MK902827, MK902837 |
| **23** | *Hypopgthalamichthys molitrix* | MK902871 |
| **24** | *Istiompax indica* | MK902861 |
| **25** | *Istiophorus platypterus* | MK902869, MK902909 |
| **26** | *Johnrandallia nigrirostris* | MK902809, MK902819 |
| **27** | *Kajikia audax* | MK902828, MK902838 |
| **28** | *Katsuwonus pelamis* | MK902847, MK902853 |
| **29** | *Kyphosus elegans* | MK902810, MK902862, MK902870 |
| **30** | *Lobotes pacificus* | MK902896 |
| **31** | *Lutjanus aratus* | MK902811, MK902863, MK902872 |
| **32** | *Lutjanus argentiventris* | MK902840, MK902849, MK902888 |
| **33** | *Lutjanus colorado* | MK902821, MK902830 |
| **34** | *Lutjanus novemfasciatus* | MK902854, MK902855, MK902901 |
| **35** | *Lutjanus peru* | MK902820, MK902829, MK902839 |
| **36** | *Lutjanus viridis* | MK902848 |
| **37** | *Microlepidotus inornatus* | MK902864 |
| **38** | *Mugil cephalus* | MK902878 |
| **39** | *Mulloidichthys dentatus* | MK902873 |
| **40** | *Mycteroperca jordani* | MK902812, MK902813, MK902822, MK902831, MK902841, MK902856, MK902865, MK902874 |
| **41** | *Mycteroperca rosacea* | MK902823, MK902832, MK902842, MK902866 |
| **42** | *Paralabrax aurogutatus* | MK902824, MK902833 |
| **43** | *Paralabrax maculatofasciatus* | MK902875 |
| **44** | *Paralabrax nebulifer* | MK902843, MK902850, MK902857 |
| **45** | *Paranthias colonus* | MK902908, MK902911 |
| **46** | *Peprilus snyderi* | MK902917 |
| **47** | *Phthanophaneron harveyi* | MK902912 |
| **48** | *Prionurus punctatus* | MK902876 |
| **49** | *Rachycentron canadum* | MK902895 |
| **50** | *Rypticus bicolor* | MK902880, MK902918 |
| **51** | *Scarus ghobban* | MK902891 |
| **52** | *Scarus perrico* | MK902897, MK902903 |
| **53** | *Scomberus sierra* | MK902814, MK902877 |
| **54** | *Sebastes macdonaldi* | MK902904 |
| **55** | *Seriola dumerili* | MK902905 |
| **56** | *Seriola lalandi* | MK902844 |
| **57** | *Seriola rivoliana* | MK902851, MK902858 |
| **58** | *Sphoeroides anulatus* | MK902882 |
| **59** | *Sphyraena ensis* | MK902890 |
| **60** | *Stegastes rectifaenum* | MK902815 |
| **61** | *Sufflamen verres* | MK902825, MK902881 |
| **62** | *Tetrapturus audax* | MK902887 |
| **63** | *Thunnus albacares* | MK902892, MK902898, MK902900 |
| **64** | *Thunnus thynnus* | MK902885 |
| **65** | *Totoaba macdonaldi* | MK902883 |
| **66** | *Xiphias gladius* | MK902916 |
| **67** | *Xystreurys liolepis* | MK902910 |

#### Table S3. Primers for library construction.

**First round primer pairs**

| **Primer Forward** | | **Adapter used for 2^nd^ PCR** | **3’ end of Illumina adapter** | | **Universal 12S Forward Primer teleo_F** | |
| --- | --- | --- | --- | --- | --- | --- |
| eDNA12SV-F | | TCGTCGGCAGCGTC | AGATGTGTATAAGAGACAG | | ACACCGCCCGTCACTCT | |
| **Primer Reverse** | | **Adapter used for 2^nd^ PCR** | **3’ end of Illumina adapter** | | **Universal 12S Reverse Primer teleo_R** | |
| eDNA12SV-R | | GTCTCGTGGGCTCGG | AGATGTGTATAAGAGACAG | | CTTCCGGTACACTTACCATG | |
| **Primers with Illumina adapters and MID for second-round PCR** | | | | | | |
| **Forward** | **Index 2** | | | **i5 MID** | | **Adapter used for 2^nd^ PCR** |
| eDNA2F-A | AATGATACGGCGACCACCGAGATCTACAC | | | GACACAGT | | TCGTCGGCAGCGTC |
| eDNA2F-B | AATGATACGGCGACCACCGAGATCTACAC | | | GCATAACG | | TCGTCGGCAGCGTC |
| eDNA2F-C | AATGATACGGCGACCACCGAGATCTACAC | | | ACAGAGGT | | TCGTCGGCAGCGTC |
| eDNA2F-D | AATGATACGGCGACCACCGAGATCTACAC | | | CCACTAAG | | TCGTCGGCAGCGTC |
| eDNA2F-E | AATGATACGGCGACCACCGAGATCTACAC | | | TGTTCCGT | | TCGTCGGCAGCGTC |
| eDNA2F-F | AATGATACGGCGACCACCGAGATCTACAC | | | GATACCTG | | TCGTCGGCAGCGTC |
| eDNA2F-G | AATGATACGGCGACCACCGAGATCTACAC | | | AGCCGTAA | | TCGTCGGCAGCGTC |
| eDNA2F-H | AATGATACGGCGACCACCGAGATCTACAC | | | CTCCTGAA | | TCGTCGGCAGCGTC |
| **Reverse** | **Index 1** | | | **i7 MID** | | **Adapter used for 2^nd^ PCR** |
| eDNA2R-01 | CAAGCAGAAGACGGCATACGAGAT | | | TCACCTAG | | GTCTCGTGGGCTCGG |
| eDNA2R-02 | CAAGCAGAAGACGGCATACGAGAT | | | CAAGTCGT | | GTCTCGTGGGCTCGG |
| eDNA2R-03 | CAAGCAGAAGACGGCATACGAGAT | | | CTGTATGC | | GTCTCGTGGGCTCGG |
| eDNA2R-04 | CAAGCAGAAGACGGCATACGAGAT | | | AGTTCGCA | | GTCTCGTGGGCTCGG |
| eDNA2R-05 | CAAGCAGAAGACGGCATACGAGAT | | | ATCGGAGA | | GTCTCGTGGGCTCGG |
| eDNA2R-06 | CAAGCAGAAGACGGCATACGAGAT | | | AAGTCCTC | | GTCTCGTGGGCTCGG |
| eDNA2R-07 | CAAGCAGAAGACGGCATACGAGAT | | | TGGATGGT | | GTCTCGTGGGCTCGG |
| eDNA2R-08 | CAAGCAGAAGACGGCATACGAGAT | | | AGGTGTTG | | GTCTCGTGGGCTCGG |
| eDNA2R-09 | CAAGCAGAAGACGGCATACGAGAT | | | GACGAACT | | GTCTCGTGGGCTCGG |
| eDNA2R-10 | CAAGCAGAAGACGGCATACGAGAT | | | GTTCTTCG | | GTCTCGTGGGCTCGG |
| eDNA2R-11 | CAAGCAGAAGACGGCATACGAGAT | | | TTCGCCAT | | GTCTCGTGGGCTCGG |
| eDNA2R-12 | CAAGCAGAAGACGGCATACGAGAT | | | CAACTCCA | | GTCTCGTGGGCTCGG |

#### S4. Details for PCR 1 and 2 in library preparation.

#### *First PCR step (PCR 1).* Each PCR1 reaction contained PCR 1x Buffer (5X Buffer Thermo), 0.2 mM dNTPs, 0.2 μM of each “teleo” F and R primers, 2 μM human blocking primer, 3% DMSO, 0.6 U Phusion polymerase (Thermo) and 1-2 μL eDNA in a total 12 μL reaction. The parameters for the thermocycling were: 98 ° C x 10min, 35 cycles of 98 ° C x 30 s, 61 ° C x 30 s, 72 ° C x 30 s, and a final extension of 72 ° C x 5 min. For all tests, negative (nuclease-free water, NEB) and positive controls (tissue-derived DNA of *Mycteroperca rosacea*) were used. Successful PCR amplifications were verified via 2% agarose gels.

***Second PCR step (PCR 2)*.** PCR amplifications contained 1x PCR Buffer (5X Buffer Thermo), 0.2 mM dNTPs, 0.2 μM F and R primers, 3% DMSO, 0.6 U Phusion (Thermo), 1-2 μL eDNA, in reactions of 12 μL. The parameters for the thermocycling were 98 ° C x 5min, 8 cycles of 98 ° C x 30 s, 61 ° C x 30 s, 72 ° C x 15 s, and a final extension of 72 ° C x 5min. We performed three PCR 2, one for each PCR 1 replicate.

**S5. Pipeline for 12S rRNA metabarcoding bioinformatic analysis (OBITOOLS, VSEARCH, SWARM, R).**

1. **Original files:**

S1_R1_001.fastq (323,473 reads), 1_S1_R2_001.fastq (323,473 reads)

2. **Merge paired-end data (archive order must be reverse R2 – forward R1).**

illuminapairedend -r 1_S1_R2_001.fastq 1_S1_R1_001.fastq > S1.fastq

3. **Keep the aligned reads with quality score > 40.**

obigrep -p ‘score>40.00’ -p ‘mode!=”joined”’ S2.fastq > S2.ali.fastq

4. **Label sample and experiment with ngsfilter.**

ngsfilter -t S2_ngsfilter.txt -u S2_unidentified.fastq S2.ali.fastq > S2.ali.assigned.fasta

5. **Obistats. To evaluate mean read length.**

obistats -c sample –mean seq_length S1.ali.assigned.fastq > S1.stat.ali.assigned.fastq

6. **Filter by length and with no Ns.**

obigrep -p ‘seq_length<66’ -p ‘seq_length>59’ -s ‘^[ACGT]+$’ S1.ali.assigned.fastq > S1.ali.filtered.fastq

7. **Concatenate all samples.**

cat *.ali.filtered.fastq > cat.filtered.fastq

8. **Dereplicate reads into unique sequences.**

obiuniq -m sample cat.ali.filtered.fastq > cat.unique.fasta

9. **Sort by abundance and change identifier of the sequence.**

obiannotate –seq-rank S1.uniq.fasta | obiannotate –set-identifier ‘”’Gulf’_%09d” % seq_rank’ cat.rank.fasta > cat.new.fasta

10. **Change format fasta to vsearch.**

Rscript owi_obifasta2vsearch -I cat.new.fasta -o cat.vsearch.fasta

11. **Remove chimaeras of all samples together with vsearch.**

vsearch –uchime_denovo cat.vsearch.fasta –sizeout –minh 0.90 –nonchimeras cat.nonchimeras.fasta –chimeras cat.chimeras.fasta –uchimeout cat.uchime_out.txt

12. **Change format from vsearch to obitools.**

Rscript owi_obifasta2vsearch -I cat.nonchimeras.fasta -o cat.nochimeras.vsearch.fasta

13. **Cluster using SWARM.**

swarm -d 2 -z -t 40 -o cat_SWARM1nc_output -s cat_SWARM1nc_stats -w cat_SWARM1nc_seeds.fasta cat.nonchimeras.fasta

14. **Recount the abundances.**

obitab -o cat.new.fasta > cat.new.tab

15. **Recount after SWARM to generate OTU table per sampling site and removes singletons after clustering**

Rscript owi_recount_swarm cat_SWARM_2_output cat.new.tab

**S6. Taxonomic list of Actinopterygii detected from UVC.**

| **Class** | **Order** | **Family** | **Genus** | **Species** |
| --- | --- | --- | --- | --- |
| Actinopterygii | Anguilliformes | Muraenidae | Gymnothorax | *Gymnothorax castaneus* |
|  |  |  | Muraena | *Muraena lentiginosa* |
|  | Aulopiformes | Synodontidae | Synodus | *Synodus lacertinus* |
|  | Beryciformes | Holocentridae | Myripristis | *Myripristis leiognathus* |
|  |  |  | Neoniphon | *Neoniphon suborbitalis* |
|  | Elopiformes | Elopidae | Elops | *Elops affinis* |
|  | Perciformes | Acanthuridae | Acanthurus | *Acanthurus nigricans* |
|  |  |  |  | *Acanthurus triostegus* |
|  |  |  |  | *Acanthurus xanthopterus* |
|  |  |  | Prionurus | *Prionurus punctatus* |
|  |  | Apogonidae | Apogon | *Apogon pacificus* |
|  |  |  |  | *Apogon retrosella* |
|  |  | Blenniidae | Ophioblennius | *Ophioblennius steindachneri* |
|  |  |  | Plagiotremus | *Plagiotremus azaleus* |
|  |  | Carangidae | Carangoides | *Carangoides orthogrammus* |
|  |  |  | Caranx | *Caranx caballus* |
|  |  |  |  | *Caranx sp* |
|  |  |  | Gnathanodon | *Gnathanodon speciosus* |
|  |  |  | Seriola | *Seriola lalandi* |
|  |  |  | Trachinotus | *Trachinotus rhodopus* |
|  |  | Chaenopsidae | Chaenopsis | *Chaenopsis alepidota* |
|  |  | Chaetodontidae | Chaetodon | *Chaetodon humeralis* |
|  |  |  | Johnrandallia | *Johnrandallia nigrirostris* |
|  |  | Cirrhitidae | Cirrhitichthys | *Cirrhitichthys oxycephalus* |
|  |  |  | Cirrhitus | *Cirrhitus rivulatus* |
|  |  | Gobiidae | Coryphopterus | *Coryphopterus urospilus* |
|  |  |  | Lythrypnus | *Lythrypnus dalli* |
|  |  | Haemulidae | Anisotremus | *Anisotremus davidsonii* |
|  |  |  |  | *Anisotremus interruptus* |
|  |  |  |  | *Anisotremus taeniatus* |
|  |  |  | Haemulon | *Haemulon maculicauda* |
|  |  |  |  | *Haemulon scudderii* |
|  |  |  |  | *Haemulon sexfasciatum* |
|  |  |  |  | *Haemulon steindachneri* |
|  |  |  | Microlepidotus | *Microlepidotus inornatus* |
|  |  | Kyphosidae | Girella | *Girella simplicidens* |
|  |  |  | Kyphosus | *Kyphosus azurea* |
|  |  |  |  | *Kyphosus elegans* |
|  |  |  |  | *Kyphosus ocyurus* |
|  |  |  |  | *Kyphosus vaigiensis* |
|  |  | Labridae | Bodianus | *Bodianus diplotaenia* |
|  |  |  | Halichoeres | *Halichoeres chierchiae* |
|  |  |  |  | *Halichoeres dispilus* |
|  |  |  |  | *Halichoeres melanotis* |
|  |  |  |  | *Halichoeres nicholsi* |
|  |  |  |  | *Halichoeres notospilus* |
|  |  |  |  | *Halichoeres semicinctus* |
|  |  |  | Semicossyphus | *Semicossyphus pulcher* |
|  |  |  | Thalassoma | *Thalassoma lucasanum* |
|  |  | Labrisomidae | Labrisomus | *Labrisomus xanti* |
|  |  |  | Malacoctenus | *Malacoctenus sp* |
|  |  | Lutjanidae | Lutjanus | *Lutjanus argentiventris* |
|  |  |  |  | *Lutjanus guttatus* |
|  |  |  |  | *Lutjanus novemfasciatus* |
|  |  |  |  | *Lutjanus viridis* |
|  |  | Mullidae | Mulloidichthys | *Mulloidichthys dentatus* |
|  |  | Pomacanthidae | Holacanthus | *Holacanthus passer* |
|  |  |  | Hoplopagrus | *Hoplopagrus guentherii* |
|  |  |  | Pomacanthus | *Pomacanthus zonipectus* |
|  |  | Pomacentridae | Abudefduf | *Abudefduf troschelii* |
|  |  |  | Chromis | *Chromis atrilobata* |
|  |  |  |  | *Chromis limbaughi* |
|  |  |  | Microspathodon | *Microspathodon bairdii* |
|  |  |  |  | *Microspathodon dorsalis* |
|  |  |  | Stegastes | *Stegastes acapulcoensis* |
|  |  |  |  | *Stegastes flavilatus* |
|  |  |  |  | *Stegastes rectifraenum* |
|  |  | Scaridae | Nicholsina | *Nicholsina denticulata* |
|  |  |  | Scarus | *Scarus compressus* |
|  |  |  |  | *Scarus ghobban* |
|  |  |  |  | *Scarus perrico* |
|  |  |  |  | *Scarus rubroviolaceus* |
|  |  | Sciaenidae | Pareques | *Pareques sp* |
|  |  | Serranidae | Alphestes | *Alphestes immaculatus* |
|  |  |  | Cephalopholis | *Cephalopholis colonus* |
|  |  |  |  | *Cephalopholis panamensis* |
|  |  |  | Epinephelus | *Epinephelus labriformis* |
|  |  |  | Mycteropeca | *Mycteroperca jordani* |
|  |  |  |  | *Mycteroperca prionura* |
|  |  |  |  | *Mycteroperca rosacea* |
|  |  |  | Paralabrax | *Paralabrax maculatofasciatus* |
|  |  |  |  | *Paralabrax sp* |
|  |  |  | Rypticus | *Rypticus bicolor* |
|  |  |  | Serranus | *Serranus psittacinus* |
|  |  | Sparidae | Calamus | *Calamus brachysomus* |
|  |  | Sphyraenidae | Sphyraena | *Sphyraena lucasana* |
|  |  | Tripterygiidae | Crocodilichthys | *Crocodilichthys gracilis* |
|  |  | Zanclidae | Zanclus | *Zanclus cornutus* |
|  | Scorpaeniformes | Scorpaenidae | Scorpaena | *Scorpaena mystes* |
|  | Sygnathiformes | Fistulariidae | Fistularia | *Fistularia commersonii* |
|  | Tetraodontiformes | Balistidae | Balistes | *Balistes polylepis* |
|  |  |  | Pseudobalistes | *Pseudobalistes naufragium* |
|  |  |  | Sufflamen | *Sufflamen verres* |
|  |  | Diodontidae | Diodon | *Diodon holocanthus* |
|  |  | Tetraodontidae | Arothron | *Arothron meleagris* |
|  |  |  | Canthigaster | *Canthigaster punctatissima* |
|  |  |  | Sphoeroides | *Sphoeroides annulatus* |

**S7. Identity % percentage estimated at intra-genus and intra-family level with the 12S rRNA barcode.**

**Species with 100% of identity**

| *Chaetodon humeralis* (Chaetodontidae) | *Anisotremus interruptus* (Haemulidae) |
| --- | --- |
| *Bodianus diplotaenia* (Labridae) | *Johnrandalia nigrirostris* (Chaetodontidae) |
| *Istiompax indica* (Istiophoridae) | *Tetrapturux audax* (Istiophoridae) |
| *Kajikia audax* (Istiophoridae) | *Tetrapturux audax* (Istiophoridae) |
| *Lutjanus novemfasciatus* (Lutjanidae) | *Hoplopagrus guentheri* (Lutjanidae) |
| *Rypticus bicolor* (Serranidae) | *Paranthias colonus* (Serranidae) |
| *Rypticus bicolor* (Serranidae) | *Epinephelus labriformis* (Serranidae) |
| *Paranthias colonus* (Serranidae) | *Epinephelus labriformis* (Serranidae) |
| *Paralabrax aurogutatus* (Serranidae) | *Paralabrax nebulifer* (Serranidae) |
| *Scarus ghobban* (Scaridae) | *Scarus perrico* (Scaridae) |

**INTRA-GENUS**

| **Genus** | **Species included** | **%** |
| --- | --- | --- |
| Cynoscion | *C. reticulatus, C. xanthulus* | 93.2 |
| Epinephelus | *E. acanthistius, E. labriformis* | 87.2 |
| Lutjanus | *L. aratus, L argentiventris, L. colorado, L. novemfasciatus, L. peru* | 85.5 – 98.6 |
| Mycteroperca | *M. rosacea, M.j ordani* | 91.9 – 93.2 |
| Paralabrax | *P. aurogutatus, P. maculatofasciatus, P.nebulifer* | 89.7 – 100 |
| Scarus | *S. ghobban, S. perrico* | 100 |
| Seriola | *S. dumerili, S. rivoliana, S. lalandi* | 94.6 – 97.3 |
| Thunnus | *T. albacares, T. thynnus* | 98.6 – 100 |

**INTRA-FAMILY**

| **Family** | **Genus included** | **Identity %** |
| --- | --- | --- |
| Scianidae | *Cynoscion, Atractoscion, Totoaba* | 90.4 – 97.3 |
| Istiophoridae | *Istiompax, Istiophorus, Kajikia, Tetrapturux* | 98 – 100 |
| Scombridae | *Katsuwomis, Thunnus, Acanthocybium* | 83.3 – 98.6 |
| Pomacentridae | *Stegastes, Abudefduf* | 57.1 |
| Haemulidae | *Anisotremus, Haemulon, Haemulopsis, Microlepidotus* | 71.2 – 91.8 |
| Chaetodontidae | *Chaetodon, Johnrandalia* | 65.4 |
| Balistidae | *Balistes, Sufflamen* | 81.1 |
| Serranidae | *Epinephelus, Mycteroperca, Paralabrax, Paranthias, Rypticus* | 57.3 – 100 |
| Carangidae | *Seriola, Gnathanodon* | 74.7 – 97.3 |
| Lutjanidae | *Lutjanus, Hoplopagrus* | 79.7 – 98.6 |

**S8. Rarefaction curves for eDNA metabarcoding sequence reads.**

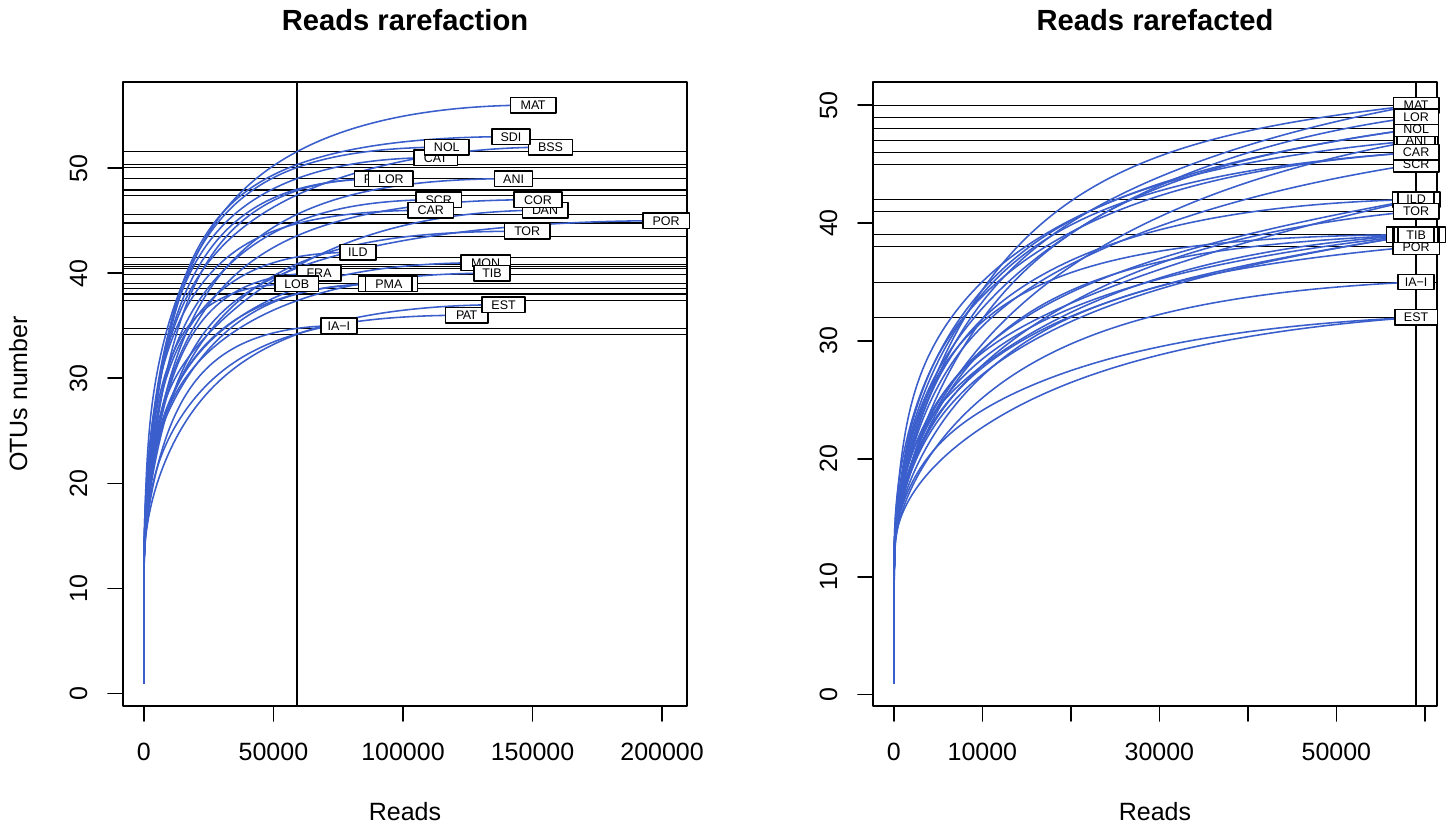

**S9. Reads counts per bioinformatic step in primary filtering.**

| **Site** | **illuminapairedend** | **obigrep** | **ngsfilter** | **obigrep** | **obiuniq** | **vsearch** | **swarm** |
| --- | --- | --- | --- | --- | --- | --- | --- |
| **POR** | 323,473 | 316,642 | 303,473 | 218,991 | 10,555 | 10,553 | 542 |
| **BSS** | 228,163 | 223,001 | 216,396 | 168,903 | 9,727 | 9,721 | 542 |
| **ANI** | 207,120 | 202,009 | 195,084 | 153,536 | 9,172 | 9,168 | 542 |
| **SDI** | 243,323 | 237,758 | 231,105 | 157,864 | 9,656 | 9,650 | 542 |
| **SCR** | 236,665 | 230,230 | 224,202 | 131,085 | 7,419 | 7,413 | 542 |
| **MAT** | 229,032 | 223,873 | 215,868 | 163,328 | 9,385 | 9,377 | 542 |
| **CAT** | 197,082 | 192,269 | 184,964 | 125,589 | 9,115 | 9,110 | 542 |
| **MON** | 198,737 | 194,233 | 188,297 | 142,806 | 8,431 | 8,428 | 542 |
| **DAN** | 231,739 | 225,811 | 217,330 | 167,751 | 9,874 | 9,869 | 542 |
| **CAR** | 248,964 | 242,700 | 220,371 | 132,021 | 8,188 | 8,182 | 542 |
| **COR** | 221,945 | 216,209 | 208,510 | 164,326 | 9,368 | 9,360 | 542 |
| **PUL** | 150,982 | 146,025 | 140,560 | 98,291 | 6,167 | 6,164 | 542 |
| **ILD** | 121,385 | 117,339 | 113,328 | 89,139 | 5,921 | 5,919 | 542 |
| **SMAR** | 170,012 | 165,282 | 159,290 | 103,654 | 6,376 | 6,374 | 542 |
| **TOR** | 214,539 | 208,221 | 206,297 | 158,662 | 8,105 | 8,102 | 542 |
| **NOL** | 198,148 | 193,252 | 184,141 | 132,260 | 7,702 | 7,694 | 542 |
| **PMA** | 212,156 | 206,831 | 196,502 | 109,218 | 6,634 | 6,631 | 542 |
| **FRA** | 121,642 | 116,636 | 112,236 | 75,776 | 6,017 | 6,015 | 542 |
| **LOR** | 206,876 | 201,315 | 193,542 | 110,516 | 7,049 | 7,046 | 542 |
| **IA-I** | 181,361 | 176,382 | 168,824 | 85,520 | 5,395 | 5,392 | 542 |
| **LOB** | 176,736 | 171,860 | 165,237 | 70,014 | 5,085 | 5,085 | 542 |
| **PAT** | 202,884 | 197,733 | 189,627 | 135,112 | 7,673 | 7,668 | 542 |
| **TIB** | 211,946 | 206,273 | 199,588 | 146,645 | 8,050 | 8,048 | 542 |
| **EST** | 206,776 | 201,297 | 191,224 | 148,228 | 8,297 | 8,287 | 542 |
| **MOCK** | 197,205 | 192,561 | 187,960 | 184,857 | 7,606 | 7,605 | 542 |
| **NEG** | 290,791 | 258,785 | 6 | 6 | 6 | 6 | 1 |

**S10. Taxonomic list of Actinopterygii detected from eDNA metabarcoding.**

| **Class** | **Order** | **Family** | **Genre** | **Species** |
| --- | --- | --- | --- | --- |
| Actinopterygii | Clupeiformes | Clupeidae | Dorosoma | *Dorosoma* sp |
|  |  |  | Sardinops | *Sardinops sagax* |
|  | Elopiformes | Elopidae | Elops | *Elops* sp |
|  | Mugilidiformes | Mugilidae | Mugil | *Mugil cephalus* |
|  | Perciformes | Acanthuridae | Acanthurus | *Acanthurus* sp |
|  |  |  |  | *Acanthurus xanthopterus* |
|  |  |  | Prionurus | *Prionurus punctatus* |
|  |  | Apogonidae | Apogon | *Apogon retrostella* |
|  |  | Balistidae | Balistes | *Balistes polylepis* |
|  |  |  |  | *Balistes* sp |
|  |  | Carangidae | Carangoides | *Carangoides* sp |
|  |  |  | Seriola | *Seriola lalandi* |
|  |  | Cirrhitidae | Cirrhitichthys | *Cirrhitichthys oxycephalus* |
|  |  | Gobiidae | Coryphopterus | *Coryphopterus uropsilus* |
|  |  | Haemulidae | Haemulon | *Haemulon sexfasciatum* |
|  |  |  | Haemulopsis | *Haemulopsis leuciscus* |
|  |  | Istiophoridae | Istiophorus | *Istiophorus platypterus* |
|  |  |  |  | OTU_01 (Istiophoridae) |
|  |  |  |  | OTU_02 (Istiophoridae) |
|  |  |  |  | OTU_03 (Istiophoridae) |
|  |  |  |  | OTU_04 (Istiophoridae) |
|  |  |  |  | OTU_05 (Istiophoridae) |
|  |  | Kyphosidae | Hermosilla | *Hermosilla azurea* |
|  |  |  | Kyphosus | *Kyphosus* sp |
|  |  |  |  | *Kyphosus elegans* |
|  |  |  | Sectator | *Sectator* sp |
|  |  | Labridae | Bodianus | *Bodianus diplotaenia* |
|  |  |  |  | *Bodianus rufus* |
|  |  |  |  | *Bodianus* sp |
|  |  |  | Semicossyphus | *Semicossyphus pulcher* |
|  |  |  | Thalassoma | *Thalassoma lucasanum* |
|  |  |  |  | OTU_06 (Labridae) |
|  |  | Lutjanidae | Lutjanus | *Lutjanus novemfasciatus* |
|  |  |  |  | *Lutjanus argentiventris* |
|  |  |  |  | *Lutjanus peru* |
|  |  |  |  | *Lutjanus* sp |
|  |  |  |  | *Lutjanus viridis* |
|  |  |  |  | OTU_07 (Lutjanidae) |
|  |  |  |  | OTU_08 (Lutjanidae) |
|  |  |  |  | OTU_09 (Lutjanidae) |
|  |  |  |  | OTU_10 (Lutjanidae) |
|  |  |  |  | OTU_11 (Lutjanidae) |
|  |  |  |  | OTU_12 (Lutjanidae) |
|  |  |  |  | OTU_13 (Lutjanidae) |
|  |  |  |  | OTU_14 (Lutjanidae) |
|  |  |  |  | OTU_15 (Lutjanidae) |
|  |  | Mullidae | Mulloidichthys | *Mulloidichthys dentatus* |
|  |  | Nomeidae | Cubiceps | *Cubiceps* sp |
|  |  | Pomacanthidae | Holacanthus | *Holacanthus* sp |
|  |  |  | Microspathodon | *Microspathodon dorsalis* |
|  |  |  | Pomacanthus | *Pomacanthus* sp |
|  |  | Pomacentridae | Abudefduf | *Abudefduf* sp |
|  |  |  |  | *Abudefduf troschelli* |
|  |  |  | Chromis | *Chromis* sp |
|  |  |  |  | *Chromis viridis* |
|  |  |  | Stegastes | *Stegastes flavilatus* |
|  |  |  |  | *Stegastes rectifraenum* |
|  |  | Scaridae | Scarus | *Scarus perrico* |
|  |  |  |  | *Scarus* sp |
|  |  |  |  | OTU_17 (Scaridae) |
|  |  |  |  | OTU_18 (Scaridae) |
|  |  |  |  | OTU_19 (Scaridae) |
|  |  |  |  | OTU_20 (Scaridae) |
|  |  | Sciaenidae | Atractosion | *Atractoscion nobilis* |
|  |  |  | Cynoscion | *Cynoscion* sp |
|  |  |  |  | *Cynoscion xanthulus* |
|  |  |  |  | OTU_21 (Sciaenidae) |
|  |  | Scombridae | Katsuwomis | *Katsuwomis pelamis* |
|  |  |  |  | *Katsuwomis* sp |
|  |  |  | Thunnus | *Thunnus albacares* |
|  |  |  |  | *Thunnus* sp |
|  |  |  |  | OTU_22 (Scombridae) |
|  |  | Serranidae | Cephalopholis | *Cephalopholis* sp |
|  |  |  | Epinephelus | *Epinephelus* sp |
|  |  |  | Hyporthodus | *Hyporthodus* sp |
|  |  |  | Mycteroperca | *Mycteroperca rosacea* |
|  |  |  |  | *Mycteroperca* sp |
|  |  |  | Paralabrax | *Paralabrax nebulifer* |
|  |  |  |  | *Paralabrax* sp |
|  |  |  | Rypticus | *Rypicus* sp |
|  |  |  |  | *Rypticus bicolor* |
|  |  |  |  | OTU_23 (Serranidae) |
|  |  |  |  | OTU_24 (Serranidae) |
|  |  |  |  | OTU_25 (Serranidae) |
|  |  |  |  | OTU_26 (Serranidae) |
|  |  |  |  | OTU_27 (Serranidae) |
|  |  |  |  | OTU_28 (Serranidae) |
|  |  | Sphyraenidae | Sphyraena | *Sphyraena ensis* |
|  |  |  |  | *Sphyraena* sp |
|  |  |  | | OTU_29 (Perciformes) |
|  |  |  | | OTU_30 (Perciformes) |
|  |  |  | | OTU_31 (Perciformes) |
|  |  |  | | OTU_32 (Perciformes) |
|  |  |  | | OTU_33 (Perciformes) |
|  |  |  | | OTU_34 (Perciformes) |
|  |  |  | | OTU_35 (Perciformes) |
|  |  |  | | OTU_36 (Perciformes) |
|  |  |  | | OTU_37 (Perciformes) |
|  |  |  | | OTU_38 (Perciformes) |
|  |  |  | | OTU_39 (Perciformes) |
|  |  |  | | OTU_40 (Perciformes) |
|  |  |  | | OTU_41 (Perciformes) |
|  |  |  | | OTU_42 (Perciformes) |
|  | Pleuronectiformes | Paralichthydae | Paralichthys | *Paralichthys* sp |
|  |  |  |  | OTU_16 (Paralichthydae) |
|  | Sygnathiformes | Fistulariidae | Fistularia | *Fistularia commersonii* |
|  | Tetraodontiformes | Diodontidae | Diodon | *Diodon liturosus* |
|  |  | | | OTU_43 (Actinopterygii) |
|  |  | | | OTU_44 (Actinopterygii) |
|  |  | | | OTU_45 (Actinopterygii) |
|  |  | | | OTU_46 (Actinopterygii) |
|  |  | | | OTU_47 (Actinopterygii) |
|  |  | | | OTU_48 (Actinopterygii) |
|  |  | | | OTU_49 (Actinopterygii) |
|  |  | | | OTU_50 (Actinopterygii) |
|  |  | | | OTU_51 (Actinopterygii) |
|  |  | | | OTU_52 (Actinopterygii) |
|  |  | | | OTU_53 (Actinopterygii) |
|  |  | | | OTU_54 (Actinopterygii) |
|  |  | | | OTU_55 (Actinopterygii) |
|  |  | | | OTU_56 (Actinopterygii) |
|  |  | | | OTU_57 (Actinopterygii) |

**S11. Species/OTU detection from UVC and eDNA metabarcoding.**

| **Specie/OTU** | **UVC** | **eDNA** |
| --- | --- | --- |
| *Abudefduf* sp |  | * |
| *Abudefduf troschelli* | * | * |
| *Acanthurus nigricans* | * |  |
| *Acanthurus* sp |  | * |
| *Acanthurus triostegus* | * |  |
| *Acanthurus xanthopterus* | * | * |
| *Alphestes immaculatus* | * |  |
| *Anisotremus davidsonii* | * |  |
| *Anisotremus interruptus* | * |  |
| *Anisotremus taeniatus* | * |  |
| *Apogon pacificus* | * |  |
| *Apogon retrostella* | * | * |
| *Arothron meleagris* | * |  |
| *Atractoscion nobilis* |  | * |
| *Balistes polylepis* | * | * |
| *Balistes* sp |  | * |
| *Bodianus diplotaenia* | * | * |
| *Bodianus rufus* |  | * |
| *Bodianus* sp |  | * |
| *Calamus brachysomus* | * |  |
| *Canthigaster punctatissima* | * |  |
| *Carangoides orthogrammus* | * |  |
| *Carangoides* sp |  | * |
| *Caranx caballus* | * |  |
| *Caranx* sp | * |  |
| *Cephalopholis panamensis* | * |  |
| *Cephalopholis* sp |  | * |
| *Chaenopsis alepidota* | * |  |
| *Chaetodon humeralis* | * |  |
| *Chromis atrilobata* | * |  |
| *Chromis limbaughi* | * |  |
| *Chromis* sp |  | * |
| *Chromis viridis* |  | * |
| *Cirrhitichthys oxycephalus* | * | * |
| *Cirrhitus rivulatus* | * |  |
| *Coryphopterus uropsilus* | * | * |
| *Crocodilichthys gracilis* | * |  |
| *Cubiceps* sp |  | * |
| *Cynoscion* sp |  | * |
| *Cynoscion xanthulus* |  | * |
| *Diodon holocanthus* | * |  |
| *Diodon liturosus* |  | * |
| *Dorosoma* sp |  | * |
| *Elops affinis* | * |  |
| *Elops* sp |  | * |
| *Epinephelus labriformis* | * |  |
| *Epinephelus* sp |  | * |
| *Fistularia commersonii* | * | * |
| *Girella simplicidens* | * |  |
| *Gnathanodon speciosus* | * |  |
| *Gymnothorax castaneus* | * |  |
| *Haemulon maculicauda* | * |  |
| *Haemulon scudderii* | * |  |
| *Haemulon sexfasciatum* | * | * |
| *Haemulon steindachneri* | * |  |
| *Haemulopsis leuciscus* |  | * |
| *Halichoeres chierchiae* | * |  |
| *Halichoeres dispilus* | * |  |
| *Halichoeres melanotis* | * |  |
| *Halichoeres nicholsi* | * |  |
| *Halichoeres notospilus* | * |  |
| *Halichoeres semicinctus* | * |  |
| *Hermosilla azurea* |  | * |
| *Holacanthus passer* | * |  |
| *Holacanthus* sp |  | * |
| *Hoplopagrus guentherii* | * |  |
| *Hyporthodus* sp |  | * |
| *Istiophorus platypterus* |  | * |
| *Johnrandallia nigrirostris* | * |  |
| *Katsuwomis pelamis* |  | * |
| *Katsuwomis* sp |  | * |
| *Kyphosus azurea* | * |  |
| *Kyphosus elegans* | * | * |
| *Kyphosus ocyurus* | * |  |
| *Kyphosus* sp |  | * |
| *Kyphosus vaigiensis* | * |  |
| *Labrisomus xanti* | * |  |
| *Lutjanus argentiventris* | * | * |
| *Lutjanus guttatus* | * |  |
| *Lutjanus novemfasciatus* | * | * |
| *Lutjanus peru* |  | * |
| *Lutjanus* sp |  | * |
| *Lutjanus viridis* | * | * |
| *Lythrypnus dalli* | * |  |
| *Malacoctenus* sp | * |  |
| *Microlepidotus inornatus* | * |  |
| *Microspathodon bairdii* | * |  |
| *Microspathodon dorsalis* | * | * |
| *Mugil cephalus* |  | * |
| *Mulloidichthys dentatus* | * | * |
| *Muraena lentiginosa* | * |  |
| *Mycteroperca jordani* | * |  |
| *Mycteroperca prionura* | * |  |
| *Mycteroperca rosacea* | * | * |
| *Mycteroperca* sp |  | * |
| *Myripristis leiognathus* | * |  |
| *Neoniphon suborbitalis* | * |  |
| *Nicholsina denticulata* | * |  |
| *Ophioblennius steindachneri* | * |  |
| OTU_01 (Istiophoridae) |  | * |
| OTU_02 (Istiophoridae) |  | * |
| OTU_03 (Istiophoridae) |  | * |
| OTU_04 (Istiophoridae) |  | * |
| OTU_05 (Istiophoridae) |  | * |
| OTU_06 (Labridae) |  | * |
| OTU_07 (Lutjanidae) |  | * |
| OTU_08 (Lutjanidae) |  | * |
| OTU_09 (Lutjanidae) |  | * |
| OTU_10 (Lutjanidae) |  | * |
| OTU_11 (Lutjanidae) |  | * |
| OTU_12 (Lutjanidae) |  | * |
| OTU_13 (Lutjanidae) |  | * |
| OTU_14 (Lutjanidae) |  | * |
| OTU_15 (Lutjanidae) |  | * |
| OTU_16 (Paralichthydae) |  | * |
| OTU_17 (Scaridae) |  | * |
| OTU_18 (Scaridae) |  | * |
| OTU_19 (Scaridae) |  | * |
| OTU_20 (Scaridae) |  | * |
| OTU_21 (Sciaenidae) |  | * |
| OTU_22 (Scombridae) |  | * |
| OTU_23 (Serranidae) |  | * |
| OTU_24 (Serranidae) |  | * |
| OTU_25 (Serranidae) |  | * |
| OTU_26 (Serranidae) |  | * |
| OTU_27 (Serranidae) |  | * |
| OTU_28 (Serranidae) |  | * |
| OTU_29 (Perciformes) |  | * |
| OTU_30 (Perciformes) |  | * |
| OTU_31 (Perciformes) |  | * |
| OTU_32 (Perciformes) |  | * |
| OTU_33 (Perciformes) |  | * |
| OTU_34 (Perciformes) |  | * |
| OTU_35 (Perciformes) |  | * |
| OTU_36 (Perciformes) |  | * |
| OTU_37 (Perciformes) |  | * |
| OTU_38 (Perciformes) |  | * |
| OTU_39 (Perciformes) |  | * |
| OTU_40 (Perciformes) |  | * |
| OTU_41 (Perciformes) |  | * |
| OTU_42 (Perciformes) |  | * |
| OTU_43 (Actinopterygii) |  | * |
| OTU_44 (Actinopterygii) |  | * |
| OTU_45 (Actinopterygii) |  | * |
| OTU_46 (Actinopterygii) |  | * |
| OTU_47 (Actinopterygii) |  | * |
| OTU_48 (Actinopterygii) |  | * |
| OTU_49 (Actinopterygii) |  | * |
| OTU_50 (Actinopterygii) |  | * |
| OTU_51 (Actinopterygii) |  | * |
| OTU_52 (Actinopterygii) |  | * |
| OTU_53 (Actinopterygii) |  | * |
| OTU_54 (Actinopterygii) |  | * |
| OTU_55 (Actinopterygii) |  | * |
| OTU_56 (Actinopterygii) |  | * |
| OTU_57 (Actinopterygii) |  | * |
| *Paralabrax maculatofasciatus* | * |  |
| *Paralabrax nebulifer* |  | * |
| *Paralabrax* sp | * | * |
| *Paralichthys* sp |  | * |
| *Paranthias colonus* | * |  |
| *Pareques* sp | * |  |
| *Plagiotremus azaleus* | * |  |
| *Pomacanthus* sp |  | * |
| *Pomacanthus zonipectus* | * |  |
| *Prionurus punctatus* | * | * |
| *Pseudobalistes naufragium* | * |  |
| *Rypicus* sp |  | * |
| *Rypticus bicolor* | * | * |
| *Sardinops sagax* |  | * |
| *Scarus compressus* | * |  |
| *Scarus ghobban* | * |  |
| *Scarus perrico* | * | * |
| *Scarus rubroviolaceus* | * |  |
| *Scarus* sp |  | * |
| *Scorpaena mystes* | * |  |
| *Sectator* sp |  | * |
| *Semicossyphus pulcher* | * | * |
| *Seriola lalandi* | * | * |
| *Serranus psittacinus* | * |  |
| *Sphoeroides annulatus* | * |  |
| *Sphyraena ensis* |  | * |
| *Sphyraena lucasana* | * |  |
| *Sphyraena* sp |  | * |
| *Stegastes acapulcoensis* | * |  |
| *Stegastes flavilatus* | * | * |
| *Stegastes rectifraenum* | * | * |
| *Sufflamen verres* | * |  |
| *Synodus lacertinus* | * |  |
| *Thalassoma lucasanum* | * | * |
| *Thunnus albacares* |  | * |
| *Thunnus* sp |  | * |
| *Trachinotus rhodopus* | * |  |
| *Zanclus cornutus* | * |  |

**Table S12. Genera detection from UVC and eDNA metabarcoding.**

| **Genus** | **UVC** | **eDNA** |
| --- | --- | --- |
| Abudefduf | * | * |
| Acanthurus | * | * |
| Alphestes | * |  |
| Anisotremus | * |  |
| Apogon | * | * |
| Arothron | * |  |
| Atractosion |  | * |
| Balistes | * | * |
| Bodianus | * | * |
| Calamus | * |  |
| Canthigaster | * |  |
| Carangoides | * | * |
| Caranx | * |  |
| Cephalopholis | * | * |
| Chaenopsis | * |  |
| Chaetodon | * |  |
| Chromis | * | * |
| Cirrhitichthys | * | * |
| Cirrhitus | * |  |
| Coryphopterus | * | * |
| Crocodilichthys | * |  |
| Cubiceps |  | * |
| Cynoscion |  | * |
| Diodon | * | * |
| Dorosoma |  | * |
| Elops | * | * |
| Epinephelus | * | * |
| Fistularia | * | * |
| Girella | * |  |
| Gnathanodon | * |  |
| Gymnothorax | * |  |
| Haemulon | * | * |
| Halichoeres | * |  |
| Hermosilla |  | * |
| Holacanthus | * | * |
| Hoplopagrus | * |  |
| Hyporthodus |  | * |
| Istiophorus |  | * |
| Johnrandallia | * |  |
| Katsuwonus |  | * |
| Kyphosus | * | * |
| Labrisomus | * |  |
| Lutjanus | * | * |
| Lythrypnus | * |  |
| Malacoctenus | * |  |
| Microlepidotus | * |  |
| Microspathodon | * | * |
| Mugil |  | * |
| Mulloidichthys | * | * |
| Muraena | * |  |
| Mycteroperca | * | * |
| Myripristis | * |  |
| Neoniphon | * |  |
| Nicholsina | * |  |
| Ophioblennius | * |  |
| Paralabrax | * | * |
| Paralichthys |  | * |
| Pareques | * |  |
| Plagiotremus | * |  |
| Pomacanthus | * | * |
| Prionurus | * | * |
| Pseudobalistes | * |  |
| Rypticus | * | * |
| Sardinops |  | * |
| Scarus | * | * |
| Scorpaena | * |  |
| Sectator |  | * |
| Semicossyphus | * | * |
| Seriola | * | * |
| Serranus | * |  |
| Sphoeroides | * |  |
| Sphyraena | * | * |
| Stegastes | * | * |
| Sufflamen | * |  |
| Synodus | * |  |
| Thalassoma | * | * |
| Thunnus |  | * |
| Trachinotus | * |  |
| Zanclus | * |  |

**Table S13. Family detection from UVC and eDNA metabarcoding.**

| **Family** | **UVC** | **eDNA** |
| --- | --- | --- |
| **Acanthuridae** | * | * |
| **Apogonidae** | * | * |
| **Balistidae** | * | * |
| **Blenniidae** | * |  |
| **Carangidae** | * | * |
| **Chaenopsidae** | * |  |
| **Chaetodontidae** | * |  |
| **Cirrhitidae** | * | * |
| **Clupeidae** |  | * |
| **Diodontidae** | * | * |
| **Elopidae** | * | * |
| **Fistulariidae** | * | * |
| **Gobiidae** | * | * |
| **Haemulidae** | * | * |
| **Holocentridae** | * |  |
| **Istiophoridae** |  | * |
| **Kyphosidae** | * | * |
| **Labridae** | * | * |
| **Labrisomidae** | * |  |
| **Lutjanidae** | * | * |
| **Mugilidae** |  | * |
| **Mullidae** | * | * |
| **Muraenidae** | * |  |
| **Nomeidae** |  | * |
| **Paralichthydae** |  | * |
| **Pomacanthidae** | * | * |
| **Pomacentridae** | * | * |
| **Scaridae** | * | * |
| **Sciaenidae** | * | * |
| **Scombridae** |  | * |
| **Scorpaenidae** | * |  |
| **Serranidae** | * | * |
| **Sparidae** | * |  |
| **Sphyraenidae** | * | * |
| **Synodontidae** | * |  |
| **Tetraodontidae** | * |  |
| **Tripterygiidae** | * |  |
| **Zanclidae** | * |  |

**Table S14. Order detection from UVC and eDNA metabarcoding.**

| **Order** | **UVC** | **eDNA** |
| --- | --- | --- |
| **Anguilliformes** | * |  |
| **Beryciformes** | * |  |
| **Clupeiformes** |  | * |
| **Elopiformes** | * | * |
| **Mugilidiformes** |  | * |
| **Myliobatiformes** | * |  |
| **Perciformes** | * | * |
| **Pleuronectiformes** |  | * |
| **Scorpaeniformes** | * |  |
| **Sygnathiformes** | * | * |
| **Tetraodontiformes** | * | * |

**Table S15. Class detection from UVC and eDNA metabarcoding.**

| **Class** | **UVC** | **eDNA** |
| --- | --- | --- |
| **Actinopterygii** | * | * |

**S16. Spearman correlation between read abundance and organism abundance and biomass, for the shared species detected with UVC and eDNA metabarcoding in 24 sites of the GOC**, * p < 0.05.

| **Species** | **Abundance** | **Biomass** |
| --- | --- | --- |
| *Abudefduf troschelii* | -0.001 | 0.073 |
| *Acanthurus xanthopterus* | 0.259 | 0.254 |
| *Apogon retrosella* | 0.007 | 0.177 |
| *Balistes polylepis* | -0.180 | -0.267 |
| *Bodianus diplotaenia* | 0.045 | 0.106 |
| *Cirrhitichthys oxycephalus* | 0.144 | -0.032 |
| *Coryphopterus urospilus* | 0.200 | 0.191 |
| *Fistularia commersonii* | -0.136 | 0.292 |
| *Haemulon sexfasciatum* | 0.180 | -0.051 |
| *Kyphosus elegans* | -0.177 | -0.064 |
| *Lutjanus argentiventris* | 0.525* | 0.415 |
| *Lutjanus novemfasciatus* | 0.270 | 0.332 |
| *Lutjanus viridis* | 0.032 | -0.017 |
| *Microspathodon dorsalis* | 0.330 | 0.298 |
| *Mulloidichthys dentatus* | 0.029 | 0 |
| *Mycteroperca rosacea* | -0.180 | -0.297 |
| *Prionurus punctatus* | -0.299 | -0.286 |
| *Rypticus bicolor* | 0.131 | 0.096 |
| *Scarus perrico* | -0.020 | 0.099 |
| *Semicossyphus pulcher* | 0.188 | -0.151 |
| *Seriola lalandi* | -0.254 | 0 |
| *Stegastes flavilatus* | 0.089 | 0.067 |
| *Stegastes rectifraenum* | 0.043 | -0.133 |
| *Thalassoma lucasanum* | 0.355 | 0.163 |
